# Complement C3- and CR3-dependent microglial clearance protects photoreceptors in retinitis pigmentosa

**DOI:** 10.1101/504647

**Authors:** Sean M. Silverman, Wenxin Ma, Xu Wang, Lian Zhao, Wai T. Wong

## Abstract

Complement activation has been implicated as an inflammatory driver of neurodegeneration in retinal and brain pathologies. However, its involvement and influence of photoreceptor degeneration in retinitis pigmentosa (RP), an inherited, largely incurable blinding disease, is unclear. We discover that markedly upregulated retinal expression of multiple complement components coincided spatiotemporally with photoreceptor degeneration in both the rd10 mouse model and in human specimens of RP, with increased complement C3 expression and activation localizing to infiltrating microglia near photoreceptors. Genetic ablation of C3 in the rd10 background resulted in accelerated structural and functional photoreceptor degeneration and altered retinal expression of inflammatory genes. These effects were phenocopied by the genetic deletion of CR3, a microglia-expressed receptor for the C3 activation product C3b, implicating an adaptive microglial-mediation mechanism involving C3-CR3 interaction. Deficiency of either C3 or CR3 resulted in deficient microglial phagocytosis of apoptotic photoreceptors in vivo, as well as increased microglial neurotoxicity to photoreceptors in vitro. These findings demonstrate a novel adaptive role for complement activation in RP that facilitates microglial clearance of apoptotic photoreceptors, without which increased proinflammatory microglial neurotoxicity ensues. These positive contributions of complement via microglial-mediated mechanisms are important in the design of immunomodulatory therapeutic approaches to neurodegeneration.

**One Sentence Summary:** Complement activation mediates adaptive neuroprotection for photoreceptors by facilitating C3-CR3 dependent microglial clearance of apoptotic cells.

## Introduction

Retinitis pigmentosa (RP), an important blinding disease worldwide, consists of a genetically heterogenous group of inherited photoreceptor degenerations involving monogenic mutations in genes expressed predominantly in photoreceptors and retinal pigment epithelial (RPE) cells (1). Although gene therapy for one specific causative mutation (RPE65) has recently gained approval (2), broadly-applicable, efficacious treatment is still unavailable for most affected patients, who typically progress to legal blindness by 40 years of age, and to severe loss of peripheral and central vision by 60 years of age (3). This unmet medical need underscores the need to understand general mechanisms that underlie the progressive loss of photoreceptors in order to discover therapies that can slow degeneration and preserve visual function. Clinical and laboratory studies of RP have revealed that in addition to the physiological changes occurring within mutation-bearing photoreceptors, the disease is characterized by non-cell autonomous neuroinflammatory changes in the retina; these changes include the presence of activated microglia (4–6) and the upregulation in inflammatory cytokine expression (7, 8), which can influence the progression of photoreceptor degeneration (9–11). However, the consequences of innate immune system activation in the retina in the context of photoreceptor degeneration, and the molecular and cellular mechanisms underlying its activation and effects, are not well understood.

Complement expression and activation constitute one important aspect of innate immune system activation in the CNS function and pathology, both in the retina and the brain. Although complement function is necessary for normal development of neural circuits through synaptic elimination (12, 13), it has been implicated as an etiologic factor in neurodegeneration (14). In the retina, polymorphisms in multiple complement-related genes have been associated with genetic risk for age-related macular degeneration (AMD)(15), a disease featuring photoreceptor apoptosis and atrophy (16), as well as complement-laden deposits and increased complement activation (17, 18). Common to both RP and AMD is the presence of activated microglia that infiltrate into the photoreceptor layer in close proximity to degenerating photoreceptors on histopathological analyses (4, 19, 20); these microglia are also thought to be sources of complement and complement regulatory factors within the retina (21–23). Although clinical investigations involving the inhibition of complement activation as a therapeutic objective have been initiated for the treatment of AMD (24), the adaptive functions vs. deleterious dysregulation of complement activation in retinal pathologies have not been clearly discerned nor have the underlying mechanisms of action linking complement and neurodegeneration been fully elucidated.

In the current study, we sought to understand the involvement of complement in photoreceptor degeneration by employing the well-characterized rd10 mouse model of RP (25) to investigate spatiotemporal patterns of complement activation occurring in the retina and to correlate these to ongoing photoreceptor degeneration and microglial activation. We investigated the function and consequences of complement activation by examining photoreceptor degeneration in rd10 mouse that were deficient for either C3, the central complement component, and CR3, the microglia-expressed receptor to iC3b, a product of C3 activation. We discovered that complement expression and activation in the retina were prominently activated by rd10-related degeneration; in particular, increased C3 expression was found primarily in infiltrating microglia, with a concurrent opsonization of degenerating photoreceptors by the C3 activation product, iC3b. The net effect of C3 activation appears to be adaptive, as genetic deletion of C3 resulted in accelerated photoreceptor degeneration, an effect that was phenocopied with the genetic deletion of CR3. We found that this adaptive complement function related to the phagocytic clearance of apoptotic photoreceptors by microglia, which helped to promote homeostasis in the degenerating retina, and to limit collateral effects that induce further non-cell autonomous photoreceptor degeneration. While dysregulated inflammatory responses involving excessive complement and microglial activation are currently thought to predominantly exacerbate retinal degeneration, our findings here conversely highlight the adaptive functions of complement and microglia that serve to promote homeostasis and limit photoreceptor loss. These factors are likely consequential in the overall understanding of the adaptive vs. deleterious effects of neuroimmune activation in the CNS and require consideration in the design and interpretation of complement-targeted therapies for retinal diseases such as RP and AMD.

## Results

### Complement expression and activation coincide spatiotemporally with photoreceptor degeneration in retinitis pigmentosa

To address the potential contribution of local complement-mediated mechanisms to photoreceptor degeneration, we first investigated how complement expression varied spatiotemporally within the retina across the period of degeneration. We performed quantitative PCR to measure retinal mRNA levels of complement components, regulatory factors, and receptors during the highly stereotypical progression of photoreceptor degeneration in the rd10 mouse. We observed that relative to postnatal day (P) 16, a time point preceding the onset of rod degeneration, retinal levels of *C3* mRNA was prominently upregulated beginning at P21, coinciding with the onset of rod degeneration, and was maintained at elevated levels (>10 fold) up to P60 (Fig. 1A, B). Increases in mRNA levels for regulatory proteins in the alternative pathway, *Cfb* and *Cfi*, were similarly elevated in magnitude and timing, while those for *Cfh* and *Cfd* were smaller and occurred later, peaking at P45 (Fig. 1C). Expression levels of complement receptors, *Cd11b*, a component of CR3, as well as *C3ar*, and *C5ar*, were elevated during rod degeneration (Fig. 1D), as were components of the classical pathway, *C1qa* and *C2* (Fig. 1E). Similar mRNA expression changes were not detected for these transcripts over the same developmental timepoints in wild type (C57BL6/J) retina during the first postnatal month (Fig. S1). These findings reveal a prominent upregulation of multiple complement components that is induced locally in the retina upon the onset and progression of rd10-related degeneration.

**Fig.1.**
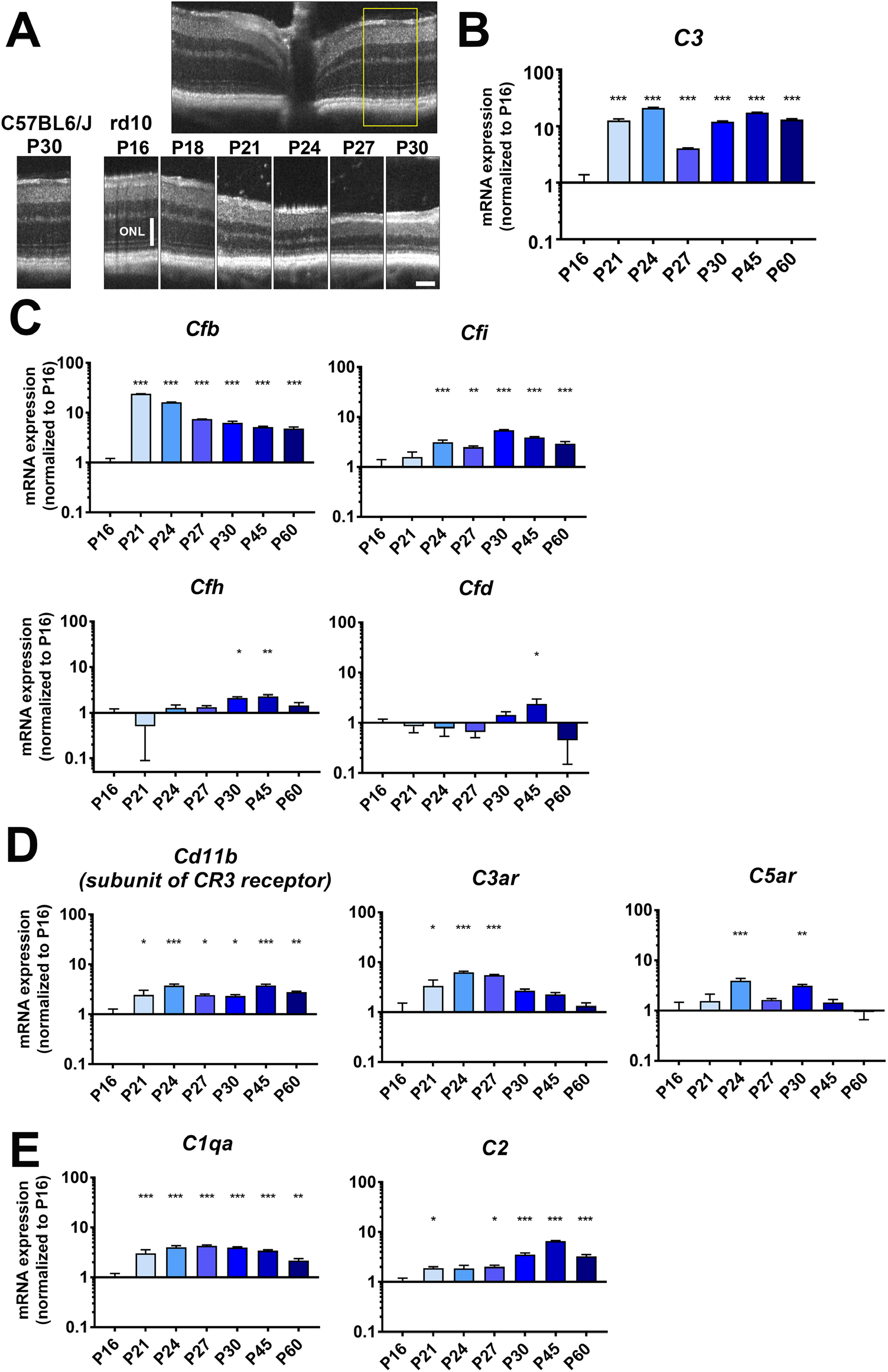
mRNA expression of complement components, regulatory factors, and receptors demonstrate prominent upregulation during photoreceptor degeneration in the rd10 mouse retina. (**A**) *In vivo* OCT of the central retina of rd10 mice demonstrated progressive atrophy of the ONL from P18, with rod degeneration being advanced by P30. Inset (yellow box) shows the retinal locus of longitudinal comparison. Scale bar = 100µm (**B**) mRNA expression levels of complement molecules in the rd10 retina from P16 to P60 were analyzed with RT-PCR. Expression of *C3* was markedly upregulated from P21 and sustained until P60. (**C**) Analysis of mRNA expression of complement regulatory factors showed early and marked upregulation for *Cfb* and *Cfi,* but not for *Cfh* or *Cfd*. (**D**) mRNA levels for receptors of complement components, including *Cd11b*, *C3ar*, and *C5ar*, as well as for components of the classical complement pathway, *C1qa* and *C2*, were also significantly increased during photoreceptor degeneration. mRNA expression levels at different time-points were normalized to levels at P16, p values for comparisons relative to levels at P16 were indicated as: *, p<0.05; **, p<0.01; ***, p<0.001; 1-way ANOVA with Dunnett’s multiple comparison test, n = 4 animals per time-point.

To discover the retinal locus of complement molecule expression, we performed fluorescent *in situ* hybridization of *C3* mRNA in rd10 retinal sections at timepoints across rod degeneration (P16 – P30). While intraretinal *C3* expression was not significantly detected at P16, it emerged prominently at P21 within the degenerating outer nuclear photoreceptor layer (ONL) and was sustained up to P30 (Fig. 2A). *C3* mRNA labeling colocalized predominantly with IBA1-immunopositive cells, which also expressed *Cx3cr1* mRNA, indicating microglia infiltrating into the ONL during rd10 degeneration (6) likely constituted the primary source of *C3* upregulation (Fig. 2B). Immunopositivity for iC3b, the product of C3 cleavage, was detected in the ONL beginning at P21, localizing to photoreceptor nuclei (Fig. 2C). Immunopositivity for CFB, the positive regulator of C3 activation, also colocalized with infiltrating IBA1+ cells in the ONL beginning at P21 (Fig. 2D). These findings indicate that microglia infiltrating the ONL at the start of rod degeneration likely contribute to C3 expression, with resulting C3 activation occurring in close proximity to degenerating photoreceptors. Upregulated *C3* expression within ONL-infiltrating microglia was similarly detected in histopathological specimens from human patients diagnosed with retinitis pigmentosa (RP), but absent in healthy, non-RP retinas (Fig. S2). This indicates *C3* upregulation as an inflammatory response in the context of inherited photoreceptor degenerations that is conserved across species.

**Fig.2.**
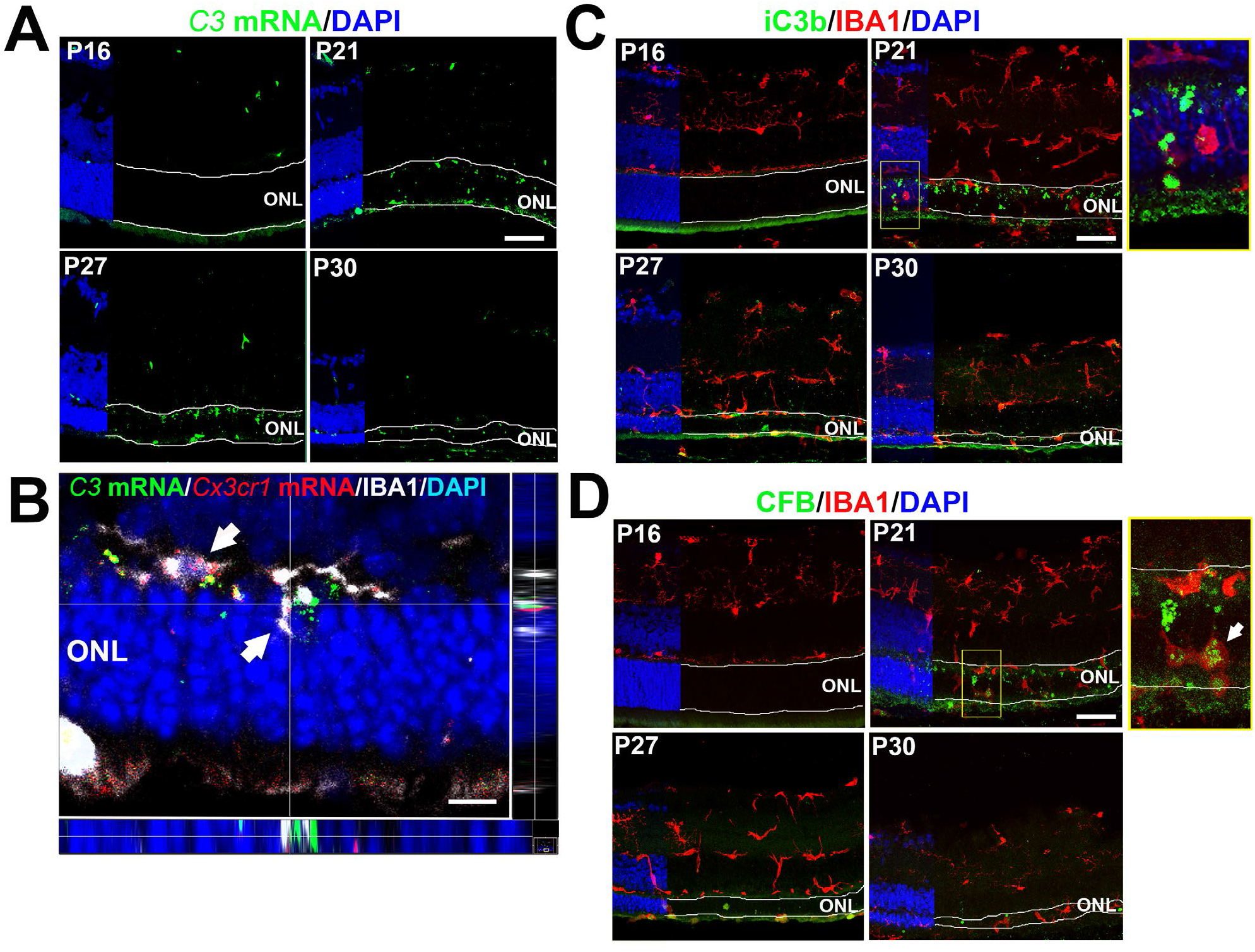
Increased complement expression and complement activation during photoreceptor degeneration in the rd10 retina are spatially localized to the degenerating ONL. (**A**) Analysis of *C3* mRNA expression by *in situ* hybridization in retinal sections from rd10 animals aged P16 – P30 demonstrated minimal labeling in P16 sections but increased labeling located specifically within the ONL (highlighted by boundary lines) in sections from P21 to P30. (**B**) High magnification analysis at P21 demonstrated that labeling for *C3* mRNA (*green*) colocalized spatially with regions of *Cx3cr1* mRNA labeling, detected with *in situ* hybridization (red), and IBA1 immunopositivity (*white*), indicating prominent *C3* mRNA upregulation within microglia infiltrating the degenerating ONL. Orthogonal views demonstrate localization of *C3* and *Cx3cr1* mRNA within the IBA1+ microglial cytoplasm, scale bar = 10µm. (**C**) Immunohistochemical analysis for iC3b (*green*), a product of C3 activation and cleavage, demonstrated minimal immunopositivity at P16, but localized labeling within the ONL beginning at P21. High magnification analysis (*inset*) showed iC3b deposition on somata of ONL cells near IBA1+ (*red*) infiltrating microglia. (**D**) Immunohistochemical analysis for CFB (*green*), a positive regulator of C3 activation, similarly demonstrates increased labeling beginning at P21, localizing in and around infiltrating IBA1+ (*red*) microglia in the ONL (*inset*). Scale bars = 50µm.

### Photoreceptor degeneration is accelerated with genetic deficiency for C3 in rd10 mice

To investigate whether C3 upregulation and activation constitute an adaptive mechanism or conversely a deleterious contribution to rd10-related photoreceptor degeneration, we crossed mice that were genetically ablated for the *C3* gene into the rd10 background, creating within individual litters three genotypes containing both, one, or neither copies of the intact *C3* gene: C3^+/+^.rd10; C3^+/-^.rd10; and C3^-/-^.rd10. Optical coherence tomographic (OCT) imaging of central retina at P16 prior to degeneration onset revealed no significant differences across all 3 genotypes in the general laminated structure of the retina or in the thicknesses of the overall retina or of the outer retinal layers (Fig. 3A, B). As rod degeneration progressed at P24 and P30, retinal thicknesses on OCT measurements were significantly decreased in animals deleted for one or both copies of the *C3* gene; mean thicknesses were least in C3^-/-^.rd10 animals and intermediate in C3^+/-^.rd10 animals. As rod outer segments degenerate at these timepoints, shallow areas of detachment of the outer retina from the underlying retinal pigment epithelial (RPE) layer emerged as previously described (26); these detachments were most prevalent in C3^-/-^.rd10 animals, followed by C3^+/-^.rd10 animals (Fig. 3A, C). Electroretinographic (ERG) assessment of dark-adapted responses (which are contributed to by both rod and cone photoreceptors) at P24 demonstrated significantly diminished a- and b- wave amplitudes in C3^+/-^.rd10 and C3^-/-^.rd10 animals, relative to C3^+/+^.rd10 animals (Fig. 3D), with the magnitude of the mean amplitudes between the genotypes ordered as: C3^+/+^.rd10 > C3^+/-^.rd10 > C3^-/-^.rd10, matching those for retinal thicknesses. At P24, cone-dominated light-adapted responses demonstrated no significant differences in a-wave amplitudes and a slight difference in b-wave amplitudes between genotypes (Fig. 3E). Later, at P35, when rod degeneration is fairly complete and cone degeneration underway (25), C3^-/-^.rd10 animals demonstrated significantly decreased cone responses in both the a- and b-wave amplitudes (Fig. S3), indicating that secondary cone degeneration following rod degeneration was also accelerated with C3 deficiency in the rd10 retina.

**Fig.3.**
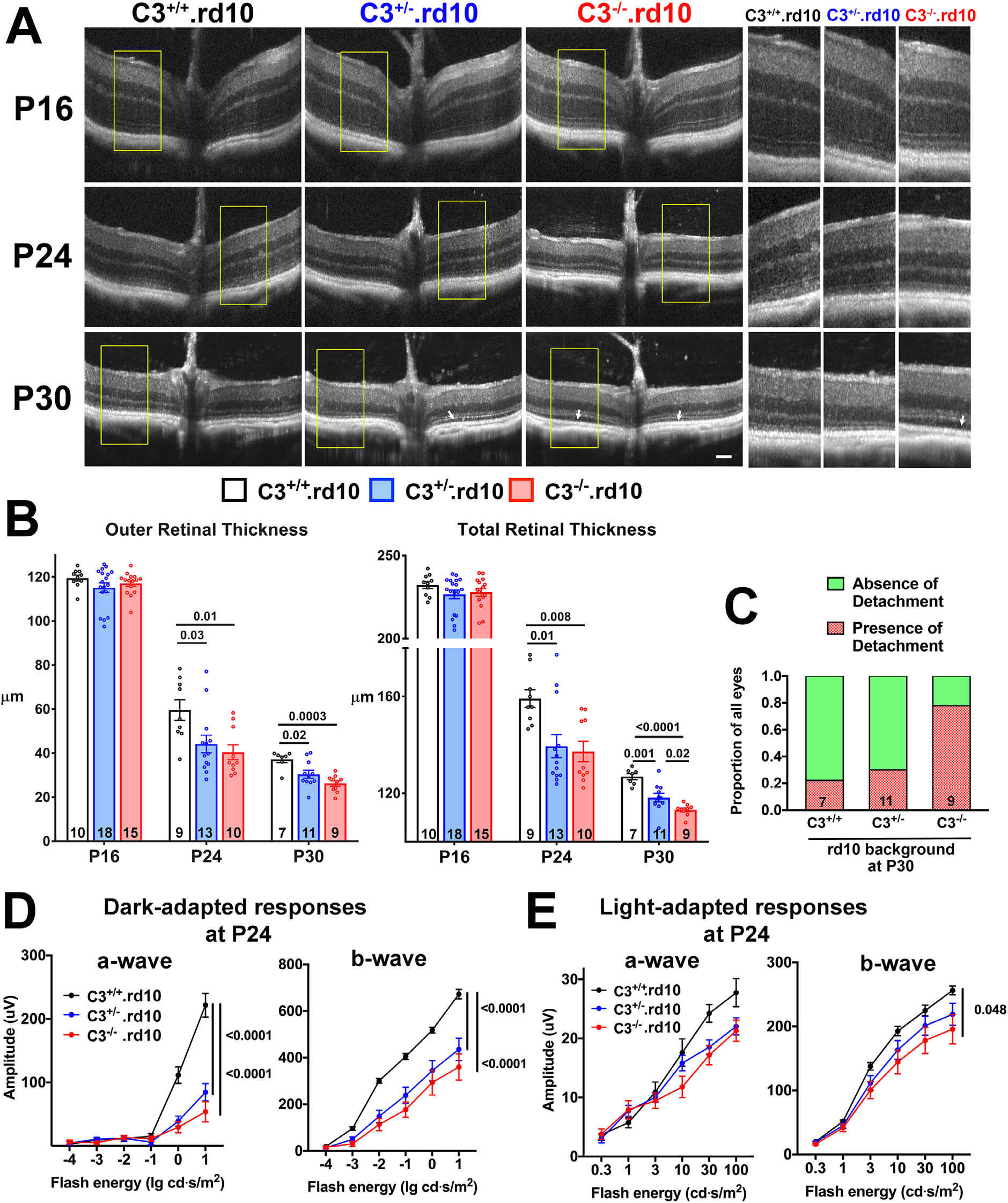
*In vivo* structural and functional evidence of accelerated photoreceptor degeneration with C3 deficiency in the rd10 mouse model. (**A**) OCT imaging was performed in mice on the rd10 in which no (C3^+/+^.rd10), one (C3^+/-^.rd10), both (C3^-/-^.rd10) copies of the *C3* gene has been genetically ablated. The range of ages of animals analyzed include time-points before (P16) and during rod degeneration (P24, P30); littermates of the 3 genotypes were analyzed and compared. Insets (*yellow boxes*) show magnified and juxtaposed images comparing corresponding retinal areas. ONL thicknesses were similar across all 3 genotypes at P16, but demonstrated more rapid decreases at P24 and P30 in C3^+/-^.rd10 and C3^-/-^.rd10 compared with C3^+/+^.rd10 retinas. Scale bar = 100µm. (**B**) Quantification of outer (from the OPL to the RPE) and total (from vitreal surface to the RPE) retinal thicknesses measured at a uniform location (350µm) radial distance from the optic nerve) demonstrated statistical similarity between the 3 genotypes at P16 but accelerated thinning in C3^+/-^.rd10 and C3^-/-^.rd10 animals (p values were derived from a 2-way ANOVA with Tukey’s multiple comparisons test, number of eyes analyzed per group provided at the bottom of each column). (**C**) The advancement of photoreceptor degeneration was accompanied by the emergence of shallow areas of retinal detachments (*white arrows* in (A)); the relative prevalence of such detachments at P30 was highest in C3^-/-^.rd10 and lowest in C3^+/+^.rd10 animals. ERG evaluation at P24 demonstrated decreased dark-adapted, rod-dominant a- and b-wave amplitudes in C3^+/-^.rd10 and C3^-/-^.rd10 relative to C3^+/+^.rd10 animals (**D**). Light-adapted, cone-mediated responses were similar in a -wave amplitudes (**E**), but slightly decreased in b-wave amplitudes for C3^-/-^.rd10 relative to C3^+/+^.rd10 animals (p values were derived from a 2- way ANOVA with Tukey’s multiple comparisons test, number of eyes analyzed were: C3^+/+^.rd10 = 12; C3^+/-^.rd10 = 11, and C3^-/-^.rd10 = 9).

Corroborating histological assessments in retinal sections revealed that ONL thickness, delineated by immunohistochemical labeling for rod-expressed rhodopsin and nuclear staining with DAPI, demonstrated significantly lower ONL thickness in C3^+/-^.rd10 and C3^-/-^.rd10 animals relative to C3^+/+^.rd10 animals, reflecting accelerated rod degeneration at both P24 and P30 (Fig. S4A, B). The densities of TUNEL+ photoreceptors in the ONL at P30 were significantly higher in C3^+/-^.rd10 and C3^-/-^.rd10 animals, reflecting increased accumulation of apoptotic photoreceptors in C3 deficiency (Fig. S4C, D).

To ascertain that C3 deficiency accelerated photoreceptor degeneration in a context related specifically to the rd10 mutation, we evaluated the structure and function of retinas from control C3^+/+^, C3^+/-^, and C3^-/-^ animals which do not possess the rd10 mutation across the same P16-P30 time period. While C3 deficiency has been previously associated with late-onset retinal degeneration in aged mice (27, 28), we did not detect C3-related retinal thinning in the first postnatal month across the same time points evaluated in rd10 mice in OCT evaluations; outer and total retinal thicknesses were similar across all 3 genotypes at P16 and P30 (Fig. S5A). ERG assessments at P24 revealed no differences in photoreceptor-dependent a-wave amplitudes, and conversely, slightly increased b-wave amplitudes in C3^+/-^ and C3^-/-^ animals relative to C3^+/+^ animals (Fig. S5B), likely reflecting decreased pruning of first-order photoreceptor synapses during retinal development (13). Taken together, these findings show that decreased C3 levels in the specific context of rd10-related changes, rather than with C3 deficiency alone, resulted in an accelerated degeneration of photoreceptors.

### Genetic deficiency for CR3, a receptor binding iC3b, accelerates rd10-related photoreceptor degeneration

We hypothesized that the mechanism underlying the adaptive function of C3 upregulation and activation in the rd10 retina involved physiological changes in retinal microglia infiltrating the photoreceptor layer. Our immunohistochemical studies had detected increased deposition of C3b/iC3b, a product of C3 activation, within the ONL in close proximity to activated infiltrating microglia, in which *Cd11b*, a component of the microglia receptor CR3, was found to be expressed. As iC3b-CR3 binding constitutes a significant mode of complement-microglia interaction enabling the clearance of opsonized debris by microglial phagocytosis (29, 30), we investigated the adaptive role of CR3-mediated microglia function by crossing mice genetically ablated for *Cd11b* (hence CR3) into the rd10 background, generating litters containing the 3 genotypes: CR3^+/+^.rd10; CR3^+/-^.rd10; and CR3^-/-^.rd10. OCT imaging demonstrated a course of photoreceptor degeneration in CR3-deficient animals that phenocopied that observed in C3-deficient animals; prior to rod degeneration (P16), no statistical differences were observed in the total retinal thickness between genotypes, however, at peak (P24) and late (P30) rod degeneration, CR3-deficient retinas demonstrated significantly greater rates of retinal thinning, with retinal thicknesses decreasing in the following order of genotypes: CR3^+/+^.rd10 > CR3^+/-^.rd10 > CR3^-/-^.rd10 (Fig. 4A). ERG assessments at P24 similarly demonstrated marked decreases in dark adapted a- and b-wave amplitudes in CR3-deficient mice relative to their CR3-sufficient littermates, while corresponding cone-mediated light-adapted responses were affected to a lesser degree (Fig. 4B). This congruence between C3- and CR3-deficiency phenotypes support the notion that C3b/iC3b signaling to retinal microglia through the CR3 receptor contributes to the complement-mediated adaptive mechanism in rd10-related photoreceptor degeneration. As found for C3 deficiency, examination of CR3^+/+^, CR3^+/-^, and CR3^-/-^ control animals lacking the rd10 mutation from P16 to P30, did not reveal evidence for structural and functional retinal degeneration (Fig. S6A, B), but conversely showed small increases in retinal thickness and b-wave amplitudes, likely due to decreased C3-CR3 mediated developmental synaptic pruning (13). These findings indicate C3-CR3 signaling as a mode of complement-microglia interaction constituting an adaptive mechanism that limits rd10-related structural and functional photoreceptor degeneration.

**Fig.4.**
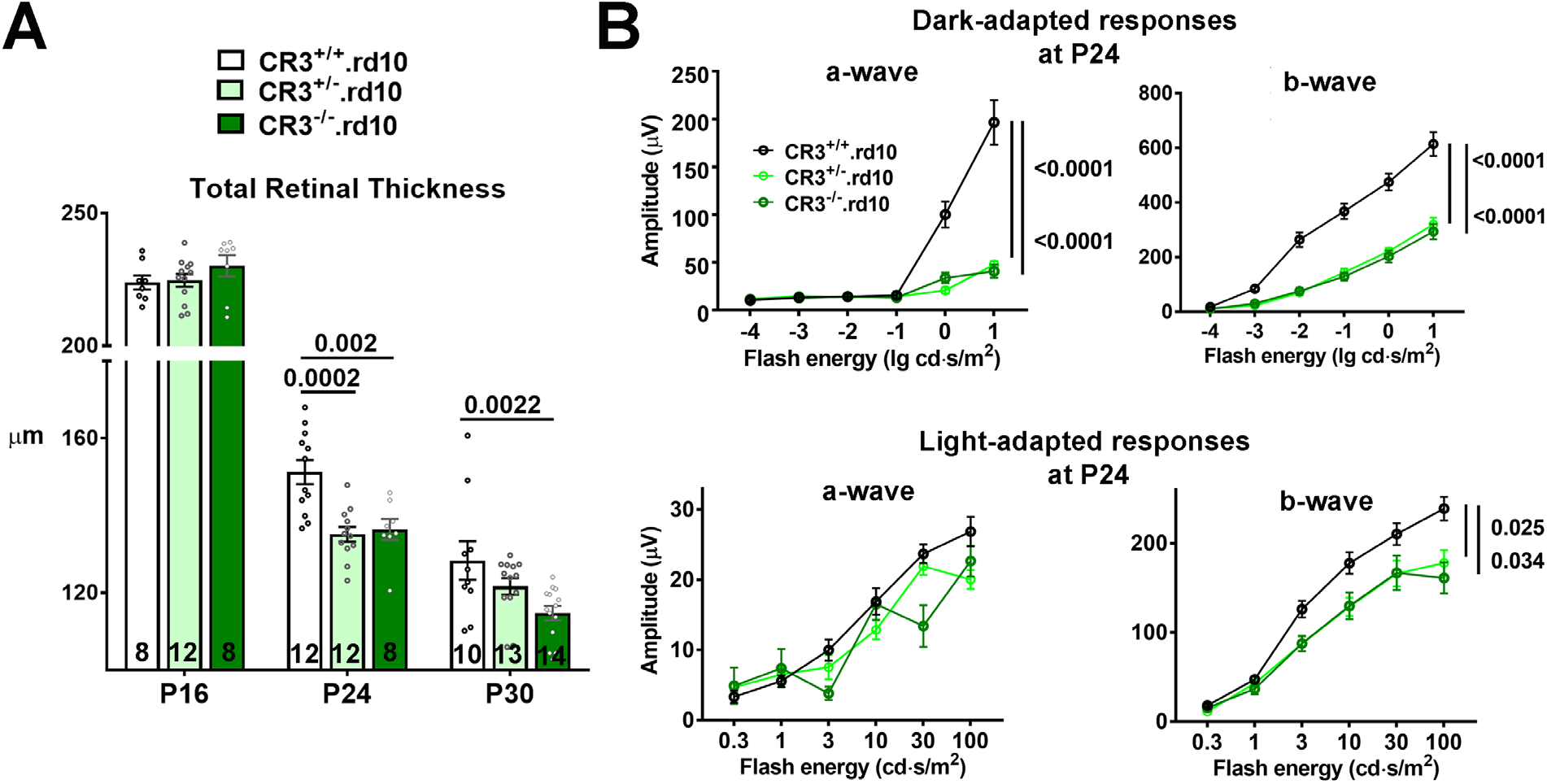
CR3 deficiency in the rd10 mouse model demonstrates an acceleration of structural and functional photoreceptor degeneration similar to that observed in C3 deficiency. (**A**) OCT imaging of the central retina was performed in mice on the rd10 background (CR3^+/+^.rd10) and compared with rd10 animals in which one (CR3^+/-^.rd10) or both (CR3^-/-^.rd10) copies of the *Cd11b* gene, a subunit of the microglial receptor, CR3, was genetically ablated. At P16, total retinal thickness was similar between all 3 genotypes. At P24 and P30, accelerated degeneration was found in CR3 ^+/-^.rd10 and CR3^-/-^.rd10 animals, relative to CR3^+/+^.rd10 animals. p values were derived from a 2-way ANOVA with Tukey’s multiple comparisons test, number of eyes analyzed in each group is provided at the bottom of each column. (**B**) ERG evaluation of retinal function at P24 demonstrated significantly and markedly lower dark-adapted, rod-dominant a- and b-wave amplitudes in CR3^+/-^.rd10 and CR3^-/-^.rd10 relative to CR3^+/+^.rd10 animals. Comparison of light-adapted, cone-mediated responses did not show significant differences in a-wave amplitudes, but b-wave amplitudes were decreased in CR3^+/-^.rd10 and CR3 ^-/-^.rd10 animals. p values were derived from a 2-way ANOVA with Tukey’s multiple comparisons test, number of eyes analyzed were: 14, 16, and 10, for CR3^+/+^.rd10, CR3^+/-^.rd10, and CR3^-/-^.rd10 animals respectively.

### C3- and CR3-deficient rd10 retinas demonstrate similarities in mRNA expression of inflammation-related genes

To further investigate the inflammatory mechanisms involving C3-CR3-mediated adaptation to photoreceptor degeneration, we compared the mRNA expression of 758 inflammatory genes in complement-sufficient rd10 retinas (C3^+/+^, CR3^+/+^.rd10) retina versus either C3- (C3^-/-^.rd10) or CR3-deficient (CR3^-/-^.rd10) retinas using the Nanostring profiling platform. At both peak (P24) and late (P30) phases of rod degeneration, unsupervised clustering of all expressed transcripts revealed greater similarity between C3^-/-^.rd10 and CR3^-/-^.rd10 retinas compared to C3^+/+^,CR3^+/+^.rd10 retinas (Fig. 5A). Identification of genes differentially expressed (DE, fold change >1.5; p value < 0.05) between (1) C3^+/+^,CR3^+/+^.rd10 vs. C3^-/-^.rd10 and (2) C3^+/+^,CR3^+/+^.rd10 vs. C3^-/-^.rd10 retinas at P24 and P30 time-points revealed that genes differentially expressed as a result of C3 deficiency overlapped considerably with those arising from CR3 deficiency, particularly at P24 (40-44% overlap)(Fig. 5B). A number of these shared DE genes were linked to microglial function and demonstrated patterns of differential expression that were similar in direction and magnitude (Fig. 5C) between C3- and CR3-deficiency. These included: *Apoe*, which has been broadly implicated in microglial responses to aging and pathological neurodegeneration (31, 32), *Lcn2*, which regulates inflammation, in part through the upregulation of microglial phagocytosis (33, 34), and *Cxcl10*, which influences microglial recruitment and debris clearance (35). Gene ontogeny analyses using IPA Ingenuity found that shared DE genes at P24 featured in canonical pathways involving interferon regulatory factor signaling, neuroinflammation signaling, and pattern recognition receptor signaling (Fig. 5D), and were associated with functions of increased neuronal and retinal degeneration, and decreased phagocyte activation and chemotaxis (Fig. 5E). These findings indicate a common molecular pattern of inflammatory changes between C3- and CR3- deficiency that appear influential to regulating the activation and phagocytic/clearance.

**Fig.5.**
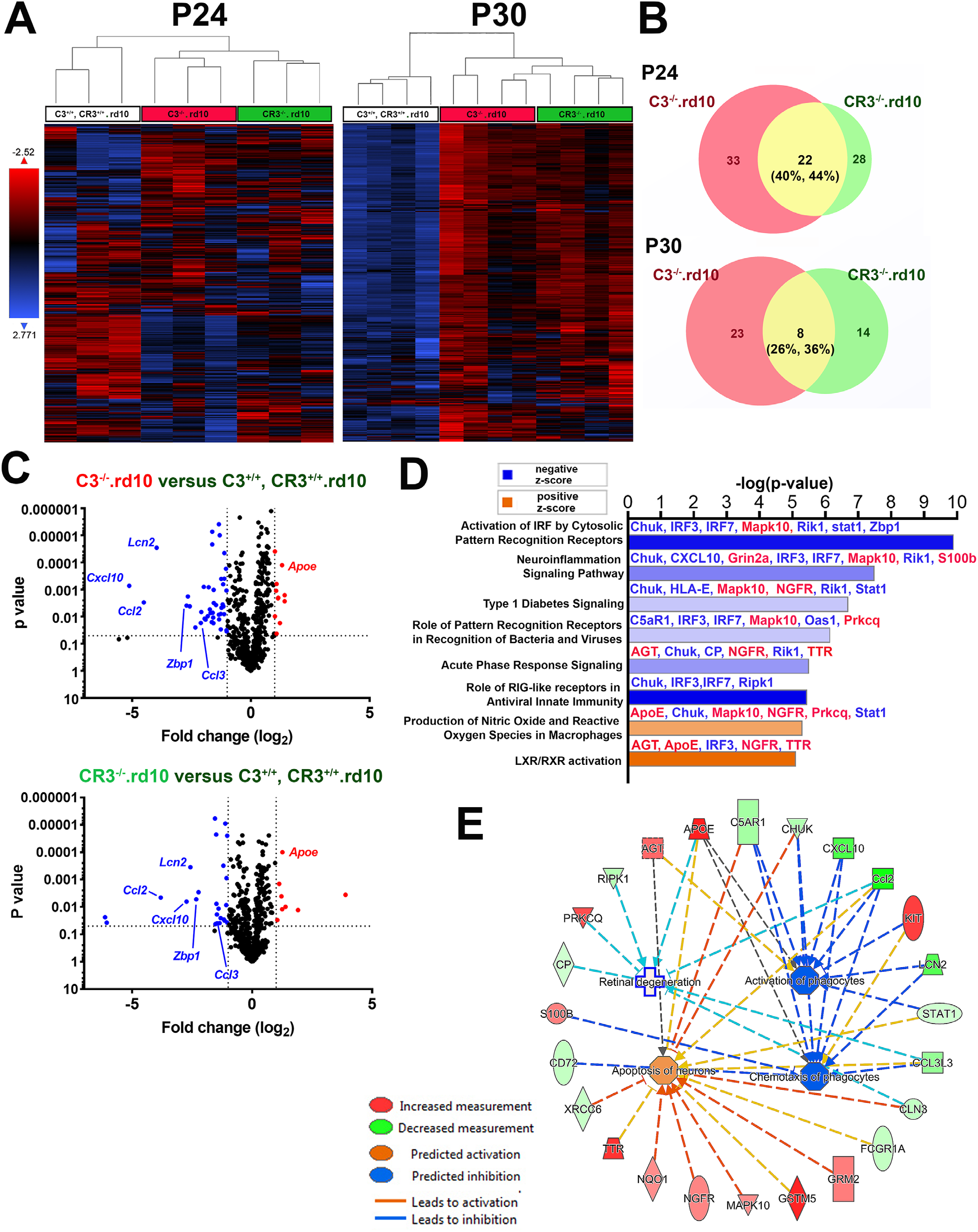
mRNA profiling of inflammatory-related genes using Nanostring reveals differentially regulated genes that are common between C3- and CR3-deficient rd10 retinas. Nanostring multiplex analysis was used to profile mRNA expression levels of 757 neuroinflammatory-related genes in retinas from the three genotypic groups at P24 and P30: (1) C3^+/+^,CR3^+/+^.rd10, (2) C3 ^-/-^.rd10, and (3) CR3^-/-^.rd10 retinas (n = 3-4 animals in each group at each age).. (**A**) Unsupervised clustering of all genes expressed above background levels demonstrated similar patterns of gene expression between C3^-/-^.rd10, and CR3^-/-^.rd10 retinas at both P24 and P30. Individual biological replicates from C3^-/-^.rd10 and CR3^-/-^.rd10 groups clustered with each other more closely than with the C3^+/+^.CR3^+/+^.rd10 group. (**B**) Genes differentially expressed (DE) (fold change ≥ 1.5; p <0.05) between C3^+/+^.CR3^+/+^.rd10 and C3 ^-/-^.rd10 retinas (*red circles*) and between C3^+/+^.CR3^+/+^.rd10 and CR3^-/-^.rd10 retinas (*green circles*) were identified at P24 and P30 and the number of DE genes in common between the two groups (*yellow segments*) counted and expressed as fractional compositions in each group). (**C**) Volcano plots highlighting the distributions of DE genes at P24 for both cross-group comparisons; annotations highlight DE genes common to both comparisons. (**D**) Listing of canonical pathways represented by common DE genes at P24 (n = 3). (**E**) Network analysis of functions attributed to common DE genes at P24.

### Microglial phagocytosis of apoptotic photoreceptors is regulated by a C3-CR3 dependent mechanism

As microglial responses in pathological situations can modulate the rate of neurodegeneration, we investigated how C3- and CR3-deficiency influences microglial morphology and physiology in the rd10 retina. We examined microglia infiltrating the ONL of flat-mounted retinas from C3-and CR3-sufficient (C3^+/+^, CR3^+/+^.rd10), C3-haploinsufficient (C3^+/-^.rd10), C3-deficient (C3^-/-^.rd10), and CR3-deficient (CR3^-/-^.rd10) animals at P24 (Fig. 6A). While the density of infiltrating microglia were similar across all 4 genotypes (Fig. 6B), the mean soma size of C3^+/+^, CR3^+/+^.rd10 microglia were significantly larger than the other 3 genotypes (Fig. 6C), corresponding to a higher mean number of phagosomes located in the soma (Fig. 6D). These differences corroborated with a greater degree of colocalization between rhodopsin immunopositivity and IBA1+ ONL microglia of C3^+/+^, CR3^+/+^.rd10 animals, indicating that decreased expression of either C3 or CR3 resulted in lower ability of infiltrating microglia to internalize degenerating photoreceptors (Fig. 6E).

**Fig.6.**
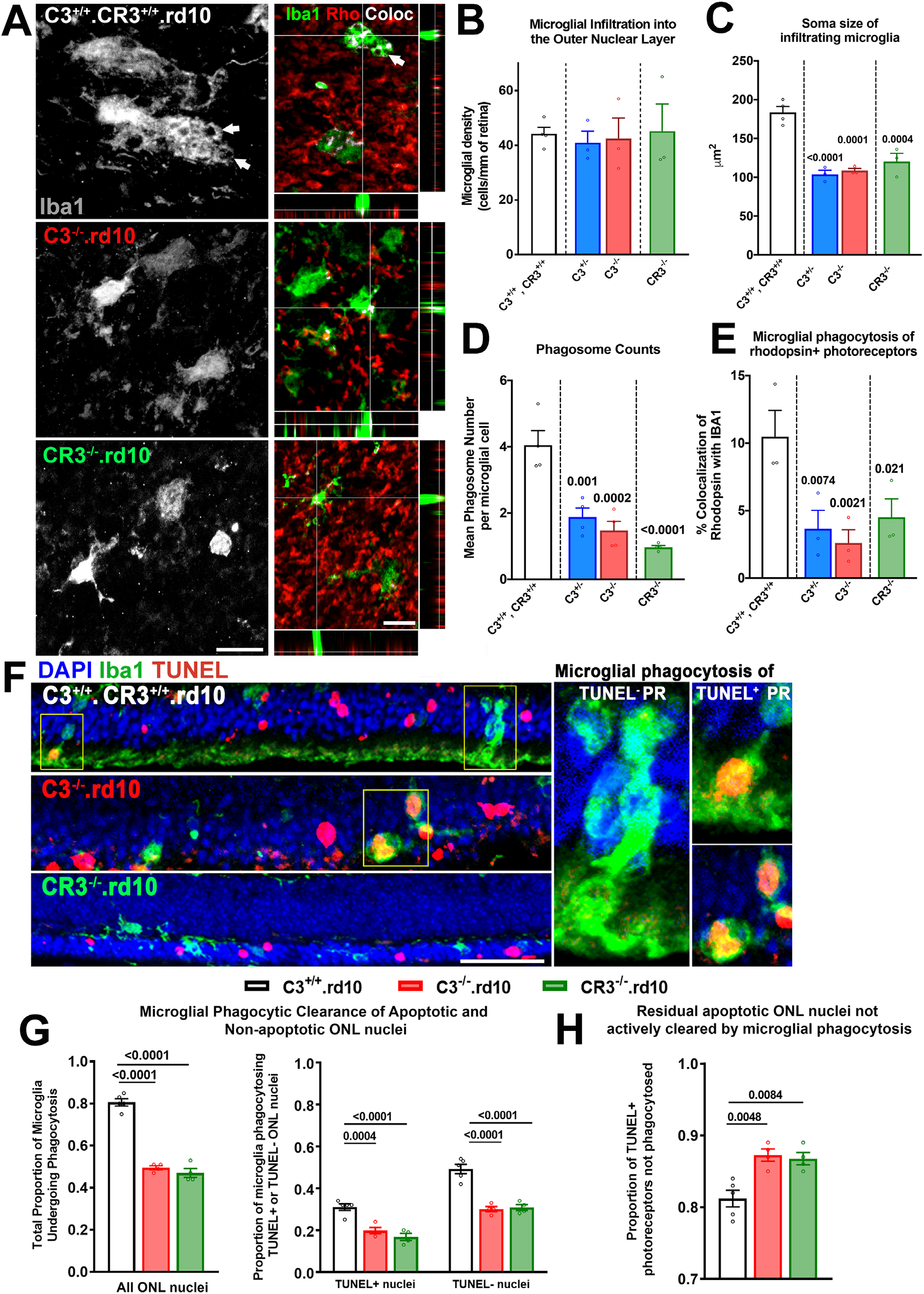
Microglial phagocytosis of apoptotic photoreceptors is decreased with C3 - and CR3-deficiency in the rd10 model of photoreceptor degeneration. **(A-E)** Immunohistochemical analysis of microglia infiltrating the ONL was performed in flat-mounted retina isolated at P24 from: rd10 retinas without C3 or CR3 deficiency (C3^+/-+^.CR3^+/+^.rd10), C3^-/-^.rd10, and CR3^-/-^.rd10 retinas. (**A**) Infiltrating IBA1+ microglia (*green*) demonstrating a deramified and amoeboid morphology were found in close contact with rhodopsin-expressing rods (*red*). Many microglia also demonstrated one or more intracellular phagosomes (*arrowheads*) that colocalized with rhodopsin immunopositivity (depicted by *white* pixels). Scale bars = 10µm. Densities of microglia infiltrating the ONL were similar between genotypes (**B**), but microglia from animals with genetic ablation of C3 or CR3 showed significantly smaller mean soma sizes (**C**), that corresponded to lower mean numbers of intracellular phagosomes per microglia (**D**), and decreased extents of rhodopsin colocalization within microglia (**E**), which reflected decreased microglial phagocytosis of rod photoreceptors. **(F-H)** Comparative analysis of phagocytosis of viable (TUNEL^-^) and apoptotic (TUNEL^+^) photoreceptors by microglia infiltrating into the ONL was performed for the 3 genotypes at P24. **(F)** Phagosomes within IBA1+ infiltrating microglia were observed to contain both TUNEL^+^ and TUNEL^-^ nuclei (*insets*). Scale bar = 50µm. (**G**) The proportions of microglia containing phagosomes were decreased in C3^-/-^.rd10 and CR3^-/-^.rd10 microglia relative to C3^+/+^.CR3^+/+^.rd10 microglia, reflecting decreased total phagocytic capacity in the absence of C3 or CR3; these differences were found for the subsets of microglia phagocytosing TUNEL+photoreceptor nuclei and phagocytosing TUNEL- photoreceptor nuclei. (**H**) The proportions of TUNEL+ nuclei left unengulfed in the ONL were increased in C3 and in CR3 deficiency, indicating a greater accumulation of uncleared apoptotic photoreceptors in these genotypes. p values indicate comparisons to the C3^+/+^.CR3^+/+^.rd10 genotype using a 1-way ANOVA with Dunnett’s multiple comparison test, n = 3-4 animals per genotypic group.

As microglia in rd10-related degeneration can phagocytose both apoptotic and non-apoptotic photoreceptors, enabling clearance of dead cells and phagoptosis of living cells respectively (6), we examined the relative phagocytosis of these targets by microglia in the contexts of C3- and CR3-deficiency (Fig. 6F). We found that the proportion of ONL microglia engaged in photoreceptor phagocytosis (containing one or more photoreceptor-containing phagosomes) was significantly decreased in C3^-/-^.rd10 and CR3^-/-^.rd10 animals compared with C3^+/+^, CR3^+/+^.rd10 animals; this measure of phagocytotic activity was decreased for both the phagocytosis of apoptotic TUNEL+ photoreceptors, as well as that of TUNEL- photoreceptors (Fig. 6G). This decreased ability of C3- and CR3-deficient microglia to phagocytose apoptotic photoreceptors also resulted in an increased proportion of TUNEL+ photoreceptors that were left uninternalized in these genotypes (Fig. 6H).

To confirm that C3-CR3 expression and interaction involving microglia directly regulate microglial phagocytic capacity, we isolated retinal microglia from C3^+/+^, CR3^+/+^.rd10, C3^-/-^.rd10, and CR3^-/-^.rd10 animals and evaluated their ability to phagocytose fluorescently-labeled bovine photoreceptor outer segments (POS) *in vitro*. In a medium supplemented with C3-depleted serum, microglia lacking either C3 or CR3 expression demonstrated a lower rate of POS internalization than microglia capable of expressing both C3 and CR3 (Fig. S7A, B). However, when exogenous C3 was added to the medium (in the form of normal serum), the proportion of microglia phagocytosing POS increased in the C3^-/-^.rd10 group to that found in the C3^+/+^, CR3^+/+^.rd10 group, while that in the CR3^-/-^.rd10 group remained unchanged. Together, these findings indicate that microglial expression and secretion of C3 can result in the opsonization of targets such as photoreceptors to promote their phagocytosis via a CR3-dependent mechanism.

### C3- and CR3-deficiency in the rd10 retina are associated with increased microglial neurotoxicity

To further explore the relationship between decreased microglial phagocytic clearance and increased photoreceptor degeneration with C3- and CR3-deficiency in the rd10 retina, we examined the ability of retinal microglia cultured from different genotypes of mice to exert neurotoxic effects on photoreceptors. Retinal microglia were isolated from P24 animals from wild type (C57BL6), C3+/+, CR3+/+.rd10, C3-/-.rd10, and CR3-/-.rd10 animals and allowed to condition serum-free media for 48 hours, which was then exposed to cultured 661W photoreceptors. While conditioned media from C3+/+, CR3+/+.rd10 microglia decreased photoreceptor viability more than microglia from wild type non-degenerating retina, conditioned media from C3- or CR3-deficient rd10 retinas exerted the largest negative impact of photoreceptor viability (Fig. 7A). Consistent with previous studies in which phagocytosis of apoptotic cells was linked with the downregulation of microglial activation and proinflammatory cytokine expression (36, 37), we found that protein levels of the cytokines TNFα, IL6, IL12, and IL33, were elevated in C3- and/or CR3-deficient rd10 retinas (Fig. 7B). These findings indicate that while increased non-clearance and accumulation of apoptotic photoreceptors owing to decreased microglial phagocytosis may directly compromise the survival of nearby photoreceptors (38), phagocytosis-deficient microglia can exert more neurotoxic effects on nearby photoreceptors, possibly via increased proinflammatory cytokine secretion. Taken together, our study examining the role of complement in photoreceptor degeneration demonstrates an adaptive, complement-dependent, mechanism that is induced upon the onset of degenerative changes that facilitates the clearance apoptotic photoreceptors by infiltrating retinal microglia expressing C3 and CR3. The inhibition of this mechanism can reduce adaptive homeostasis to retinal pathology, inducing further deleterious alterations in the immune environment of the retina, that results in an acceleration of structural and functional degeneration.

**Fig.7.**
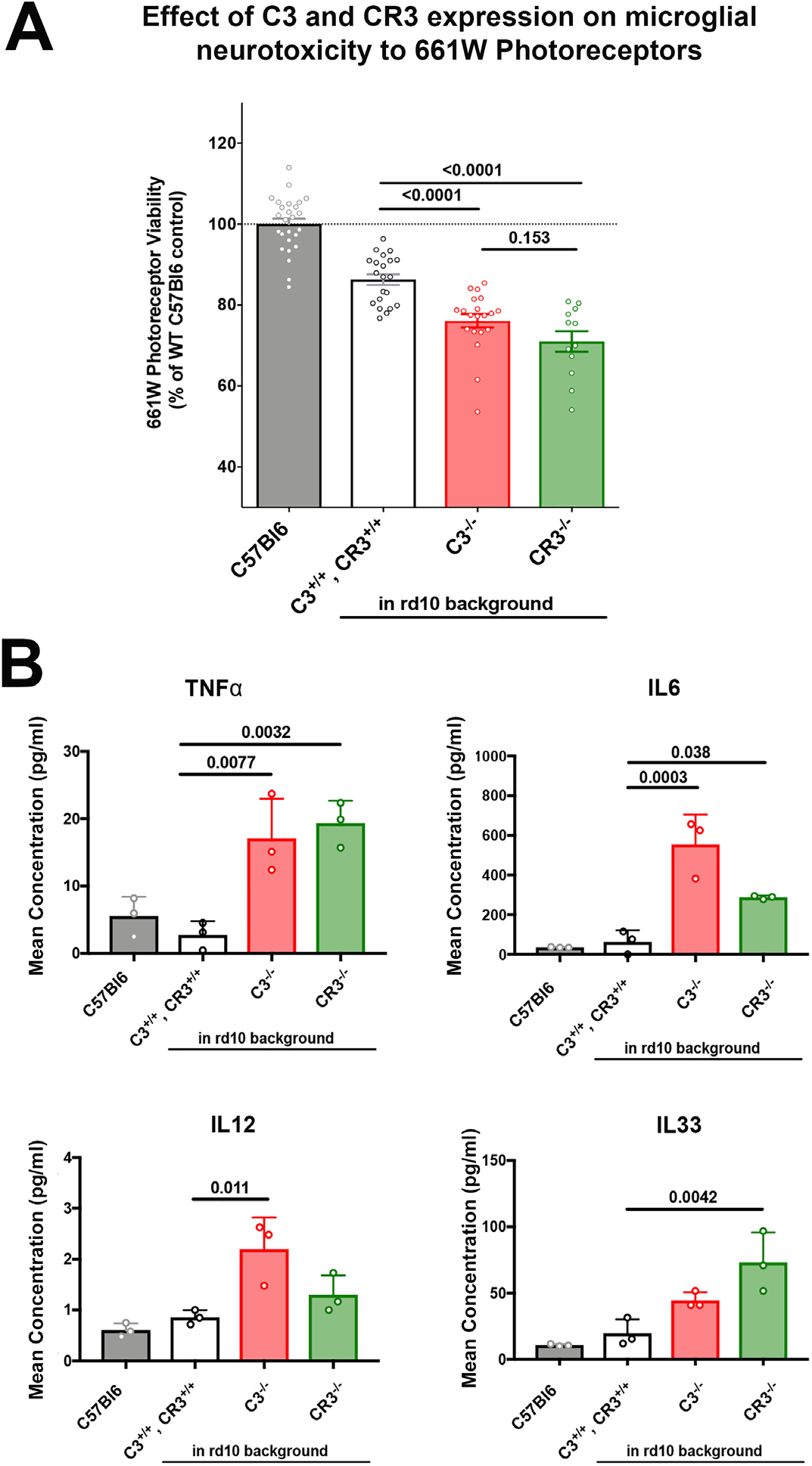
Decreased microglial phagocytic capacity from C3-CR3 deficiency is correlated with increased microglial neurotoxicity to 661W photoreceptors. (**A**) Retinal microglia were cultured from the following genotypes of mice at P24: (1) C57Bl6 wild type, (2) C3^+/+^,CR3^+/+^.rd10, (3) C3^-/-^.rd10, and (4) CR3^-/-^.rd10, and allowed to condition serum-free media for 48 hours. Direct neurotoxicity of microglia was assessed *in vitro* by exposing microglia-conditioned media to 661W photoreceptors for 12 h and evaluating photoreceptor survival using a MTT assay. Microglia from degenerating rd10 retinas of all 3 genotypes was associated with a neurotoxic effect than microglia from C57BL6 retinas. Phagocytosis-deficient microglia from C3^-/-^.rd10, and CR3^-/-^.rd10 retinas in addition were associated with significantly greater neurotoxicity than C3^+/+^,CR3^+/+^.rd10 retinas. **(B)** Increased neurotoxicity in C3^-/-^.rd10, and CR3^-/-^.rd10 microglia was associated with increased secretion of proinflammatory cytokines TNFα, IL6, IL12, and IL33 in their conditioned media, compared with C57Bl6 wild type and C3^+/+^,CR3^+/+^.rd10 microglia. p values were derived from a 1-way ANOVA with Tukey’s multiple comparisons test, n = 14 and 3 replicates per condition for the neurotoxicity assay (A) and cytokine level assay respectively.

## Discussion

We discover here that RP-related photoreceptor degeneration, which arises from a cell-autonomous photoreceptor gene mutation, induces a marked, non-cell autonomous, neuroinflammatory response that involves microglial activation and local upregulation of complement expression and activation in the photoreceptor layer. This response is spatiotemporally coincident with photoreceptor degeneration and constitutes an adaptive response aimed at restoring homeostasis by microglial phagocytic clearance of apoptotic photoreceptors, limiting the deleterious sequelae of cell death in the retina. The mechanism underlying this response, comprising microglial secretion of C3 and microglial responses via the receptor CR3, provides insight into how the innate immune system copes with the disruptive influence of intraparenchymal neuronal death and presents opportunities for immunomodulatory therapy to optimize preservation of photoreceptor structure and function in RP and other neurodegenerative retinal diseases.

Studies examining multiple CNS regions have documented complement upregulation and expression upon neurodegeneration onset in the brain (29) and the retina (36), indicating that these constitute a conserved response by the innate immune system to neuronal injury. However, corresponding to the diversity of complement-related effector functions, the net impact of complement activation on neural outcomes has been quite varied, with complement inhibition by genetic or pharmacological means producing both positive and negative outcomes depending on the context. In the majority of studies, complement activation has been associated with exacerbated neurodegeneration; genetic and/or pharmacological inhibition of complement components, such as C1q (37, 38), C3 (22, 39–41), CFB and CFD (42), have decreased retinal neurodegeneration, while genetic ablation complement components in models of Alzheimer’s disease (AD) (29, 43) and frontotemporal dementia (44), have decreased neural loss in the brain. These findings indicate that complement-mediated responses triggered in these injury contexts are likely excessive and/or inappropriately regulated and culminate in worsened overall synaptic degeneration and neuronal death. There is however emerging evidence that in other contexts complement activation may conversely serve adaptive immune functions. In the retina, absent any induced injury, long-term genetic ablation of complement molecules have resulted in a slow augmented deterioration of structure and function that gradually becomes evident in aged animals (27, 28, 45, 46), indicating that constitutive complement function may be required for long-term tissue homeostasis. In the DBA/2J mouse model of glaucoma, genetic and pharmacological inhibition of C3 increased retinal ganglion cell degeneration (47), while in the amyloid precursor protein (APP) transgenic AD mouse model, genetic C3 ablation worsened total Aβ and fibrillar amyloid plaque burden in the brain, altered microglial activation, and increased hippocampal neuronal degeneration (48). These examples indicate that complement can exert positive adaptive effects in CNS pathologies, although the underlying cellular and molecular mechanisms in these situations are not well understood.

We discover here that part of the adaptive complement response to rod photoreceptor degeneration resulting from the rod-specific *Pde6b* mutation in the rd10 mouse model involves: (1) the increased expression of C3, primarily by microglia infiltrating the photoreceptor layer at the outset of rod degeneration, (2) the opsonization of photoreceptors by iC3b, the activation product of C3, and (3) the phagocytic clearance of photoreceptors by infiltrating microglia via the CR3 receptor (Fig. S8). In the retina, while the expression of C3 has been attributed to astrocytes (47) and RPE cells (49), we find in the rd10 retina, as well as in human histopathological specimens of RP, that activated infiltrating microglia in the outer retina comprise the main source of C3 expression, similar to that found in a model of light-induced photoreceptor injury (36). The increased iC3b deposition in the ONL upon the onset of photoreceptor degeneration, elevated retinal expression of *Cd11b*, a component of the CR3, and the infiltration of CR3-expressing microglia, point to increased microglia-photoreceptor interactions via iC3b-CR3 binding (30, 50) as part of an adaptive complement response. C3-CR3 interaction has been previously demonstrated as a molecular mechanism employed by microglia to eliminate supernumerary synapses during normal brain development (13), and clear synaptic debris following injury-induced synaptic degeneration (51), but which may drive maladaptive phagocytosis of synapses in certain CNS pathologies, resulting in functional losses (29, 52). Additionally, iC3b-CR3 function in microglia has also been shown to mediate beneficial clearance of apoptotic neurons (53) and extracellular debris (54, 55) following neural injury via phagocytosis. In the context of photoreceptor degeneration, the positive vs. negative contributions of phagocytosis in microglia have also been mixed. Microglia have been shown to accelerate degeneration by phagocytosing stressed but still living photoreceptors in a process termed phagoptosis (56); inhibition of this form of phagocytosis in the rd10 model by the blockade of the vitronectin receptor resulted in slowed degeneration (6), while its augmentation in CX3CR1 deficiency accelerated loss of photoreceptor structure and function (11). Conversely, in a model of retinal detachment, retinal microglia appear to support photoreceptor survival; when autofluorescent microglia, which presumably had phagocytosed photoreceptors, were pharmacologically depleted, degeneration was increased (57). We discover here in the rd10 retina that both C3 and CR3 participate in facilitating microglial phagocytosis of apoptotic photoreceptors; when either of these factors are deficient, decreased microglial phagocytosis was accompanied by increased accumulation of apoptotic cells, increased proinflammatory cytokine expression, and accelerated degeneration. These current and previous findings indicate that microglia employ a variety of molecular mechanisms to contribute to a balance between the adaptive clearance of apoptotic cells and the maladaptive phagoptosis of stressed but still living photoreceptors.

We found here that apoptotic cell accumulation, occurring here from a deficiency in C3-CR3 mediated microglial clearance, was associated with compromised survival of the remaining photoreceptors. The mechanism for this deleterious influence may arise from apoptotic photoreceptors releasing TNFα to induce non-cell-autonomous apoptosis in adjacent cells (58, 59), and/or increased proinflammatory effects of greater microglial activation, which may be triggered by signals released from the lysis of uncleared cells (60) and decreased downregulatory effects on microglial activation that typically occur after the successful phagocytosis of apoptotic cells (61). We found support for a microglia-mediated deleterious influence as phagocytosis-deficient microglia cultured from C3- or CR3-ablated retinas demonstrated greater *in vitro* neurotoxicity to photoreceptors, an effect that was also associated with increased retinal levels of proinflammatory cytokines, including TNFα and IL6. This altered inflammatory milieu in the rd10 ONL may carry over beyond the period of rod death to secondarily impact cone death as was observed here, underscoring a broad contribution to complement-mediated apoptotic cell clearance to promoting long-term homeostasis in the degenerating retina.

One of the limitations of our study is that relatively early onset of degeneration in the rd10 retina had largely precluded the use of “cell-fate” mapping to differentiate endogenous microglia from potentially infiltrating monocytes, as we had previously done (62). Despite this, we have used the term “microglia” to refer to the IBA1+ myeloid cells infiltrating the ONL and engaging in C3 upregulation and phagocytosis of photoreceptors. While studies of other genetic mouse models of photoreceptor degeneration have described the infiltration of CCR2+ monocytes into the subretinal space (20, 63), we had previously found that in the rd10 retina, these infiltrating macrophages appeared to be largely confined to the subretinal space and did not engage photoreceptors via dynamic phagocytosis as observed for CCR2-negative microglia within the ONL (6). In other models of photoreceptor degeneration involving more acute injury, endogenous microglia, as tracked by cell-fate mapping, were found to infiltrate from the inner to the outer retina, while monocyte-derived macrophages enter into the inner retina via the retinal circulation; however these cells were not found substantially in the ONL to contribute to photoreceptor phagocytosis (62, 64). While we are unable to completely rule out possible contributions from monocyte-derived macrophages to phagocytic clearance, overall available evidence indicates endogenous infiltrating microglia to be the predominant cell type involved.

As a result of studies associating increased complement activation with retinal degeneration and implicating complement-related genes as conferring genetic risks of age-related macular degeneration (AMD) (15, 65), there is much current interest in strategies to inhibit complement activation as a treatment for degenerative retinal diseases (66). While interventions targeting C3 and complement regulatory factors to dampen complement activation in the retina (24, 67) have entered clinical investigation, the underlying molecular and cellular mechanisms involving complement in pathological retina are still obscure. Our findings in this study are useful here as they help define the adaptive contributions of complement in the diseased retina and highlight the possibility that therapeutic downregulation of complement activation may entail decreased apoptotic cell clearance thereby inducing deleterious effects. It is not clear what level of complement suppression may balance positive vs. negative contributions in each pathological context; we found in our study that even in rd10 animals heterozygous for genetic deletions of C3 or CR3 demonstrated significantly accelerated degeneration, resembling clinical examples of pathologic haploinsufficiency for complement factors documented for atypical haemolytic uraemic syndrome (68) or hereditary angioedema (69). Together, our findings highlight complexities in the multiple adaptive vs. deleterious roles of complement in retinal diseases that may require interventions to exert calibrated modulation, rather than broad and complete inhibition, for optimal therapeutic effects. Preclinical studies considering the choice of complement target and the level of complement target inhibition should assay for the effects of proposed interventions on complement- and microglia-mediated homeostatic functions to anticipate overall clinical impact and efficacy.

## Materials and Methods

### Study design

The objective of our study was to determine the presence of complement activation in RP retinas and its influence on photoreceptor degeneration by studying a well-characterized RP mouse model (rd10) and human histopathological RP specimens. We used a combination of rtPCR, *in situ* hybridization, and immunohistochemical techniques to quantitate and localize mRNA and protein expression of complement components across the period of degeneration in the RP context. Because we found the central complement component C3 and its activation product, iC3b, to be prominently upregulated in the locus of degeneration in association with infiltrating microglia, we examined the roles of C3 and its microglia-expressed receptor CR3, in degeneration by examining rd10 animals that had been crossed into C3- or CR3-deficient backgrounds, using histological techniques and *in vivo* imaging and electrophysiological assays. Changes in microglial physiology, as a result of C3- or CR3-deficiency, were also studied using *in vitro* assays of phagocytosis or neurotoxicity. Evaluation of animals in litters containing mixed genotypes were performed in a blinded manner. Biological replicate numbers are stated with each corresponding result along with the specific statistical tests used. Experiments were conducted according to protocols approved by a local Institutional Animal Care and Use Committee and adhered to the Association for Research in Vision and Ophthalmology (ARVO) Statement animal use in ophthalmic and vision research.

### Experimental animals

Experiments were conducted according to protocols approved by a local Institutional Animal Care and Use Committee and adhered to the Association for Research in Vision and Ophthalmology (ARVO) Statement animal use in ophthalmic and vision research. The following strains of mice were obtained from The Jackson Laboratory (Bar Harbor, ME): mice homozygous for the *Pde6b^rd10^* (*rd10*) mutation (#004297), mice homozygous for complement C3 mutation (#029661, C3^-/-^), and mice homozygous for the *Itgam^tm1Myd^* targeted mutation (#003991, CR3^-/-^). Animals were housed in a National Institutes of Health animal facility under 12-h light/dark cycle with food *ad libitum*. To assess complement deficiency in rd10 animals, mice were crossed with C3^-/-^ or CR3^-/-^ animals to generate animals heterozygous for these mutations in the rd10 background, which were interbred to yield animals of all three genotypes (e.g. C3^+/+^.rd10, C3^+/-^.rd10, C3^-/-^.rd10) in the same litter. Similarly, C3^-/-^ and CR3^-/-^ were bred with wild type C57BL6/J animals to generate all three genotypes in the same litter for age-matched comparisons. All experimental animals were genotyped by gene sequencing to confirm the absence of the rd8 mutation (70). Both male and female mice in the age range of postnatal days 16 to 35 were used as specified in individual experiments.

### Human Eye Tissue

Adult human eyes with clinically diagnosed RP were obtained from the donor programs of the Foundation Fighting Blindness (Columbia, MD). Retinal tissue was obtained from the following archived specimens: FFB-316 and FFB-340. Healthy control eyes without diagnoses of retinal disease were collected by Lions Eye Bank (Omaha, NE), Georgia Eye Bank (Atlanta, GA), Southern Eye Bank (Metairie, LA), and Old Dominion Eye Foundation (North Chesterfield, VA), and distributed through NDRI (Philadelphia, PA). Eyes were fixed with 2–4% paraformaldehyde, the retinas were dissected out and sectioned into 12□m-thick cryostat sections. Eye tissue was collected under applicable regulations and guidelines with proper consent, protection of human subjects, and donor confidentiality.

### Detection and localization of mRNA in retinal sections using multiplex in situ hybridization

For in situ hybridization of mRNA probes, frozen retinal tissues from murine and human eyes were sectioned at 12µm. Resulting sections were heated at 65□C for 1 h, immersed in 4% paraformaldehyde for 15 minutes, and then in RNAscope^®^ Protease III for 30 minutes at 40□C. In situ detection of mouse C3 (Mm-C3, #417841), mouse Cx3cr1 (Mm-Cx3cr1, #314221), and human C3 (Hs-C3, #430701) was performed using the RNAscope^®^ Multiplex Fluorescent Reagent Kit v2 (Advanced Cell Diagnostics) according to the manufacturer’s specifications. Fluorescent signal amplification was performed with TSA^®^ Plus fluorescein or Cy5 (1:1500, NEL760001KT, PerkinElmer).

### In vivo optical coherence tomographic (OCT) imaging of mouse retina

Mice were anesthetized with intraperitoneal ketamine (90 mg/kg) and xylazine (8 mg/kg) and their pupils were dilated. Retinal structure was assessed using an OCT imaging system (Bioptigen; InVivoVue Software). Volume scans consisting of 100 horizontal sequential B-scans (consisting of 1000 A-scans each), spanning an *en-face* retinal area of 1.4mm by 1.4mm centered on the optic nerve were captured. Retinal thickness measurements were performed in each retinal quadrant at a radial distance of 0.7mm from the optic nerve using the manufacturer’s software (Bioptigen; Diver) and averaged for each eye. Total retinal thickness, defined as the distance from the nerve fiber layer to the RPE, and outer retinal thickness, defined as the distance from the outer plexiform layer to the inner surface of the RPE, were measured from OCT images after manual retinal segmentation. The presence of shallow retinal detachments was manually evaluated in the entire scan field and scored as present or absent.

### Statistical analysis

Statistical analyses were performed using Prism 7.0d (GraphPad). For comparisons involving two data columns, t tests (paired or unpaired) or nonparametric tests (Mann–Whitney) were used, depending on whether the data followed a Gaussian distribution as determined by normality tests. A normality test (D’Agostino and Pearson) was used to analyze the distribution of all datasets. For comparisons involving three or more data columns, a one-way ANOVA (with Dunnett’s multiple-comparisons test) was used if the data followed a Gaussian distribution and a nonparametric Kruskal–Wallis test (withDunn’s multiple-comparisons test) was used if it did not. Datasets from different genotypes were compared using a two-way ANOVA.

## List of Supplementary Materials

**Supplementary Materials and Methods**

**Supplementary Figures**

Fig. S1. Complement components, regulatory factors, and receptors are not markedly increased in expression in wild type retina across the ages of P16 to P30.

Fig. S2. *C3* mRNA expression is upregulated in the ONL of histopathological retinal specimens of human retinitis pigmentosa (RP) and is localized to infiltrating microglia.

Fig. S3. C3 deficiency in the rd10 mouse model results in accelerated functional cone degeneration.

Fig. S4. Histological evidence of accelerated rod photoreceptor degeneration in C3 deficiency in the rd10 mouse model.

Fig. S5. Effect of C3 genetic deficiency in the absence of the rd10 mutation on retinal thickness and ERG function.

Fig. S6. Effect of CR3 genetic deficiency in the absence of the rd10 mutation on retinal thickness and ERG function.

Fig. S7. *In vitro* C3 opsonization of photoreceptor outer segments mediates their phagocytosis by microglia via a C3-CR3 dependent mechanism.

Fig. S8. Schematic depicting a C3-CR3-dependent mechanism of microglial phagocytic clearance of apoptotic photoreceptors in the degenerating rd10 retina.

## Acknowledgments

The authors gratefully acknowledge Drs. Joe Hollyfield and Vera Bonilha (Cleveland Clinic Cole Eye Institute Department of Ophthalmic Research) for providing access to the Foundation Fighting Blindness human RP eye collection, and Dr. Robert Fariss for the gift of human control retinal tissues.

## Funding

This study is supported by funds from the National Eye Institute Intramural Research Program (to W.T.W).

## Author contributions

SMS and WTW designed the study. SMS, WM, LZ, and XW developed the methodologies and acquired *in vivo* data. SMS and XW performed cell culture and *in vitro* assays. SMS, XW, and WTW analyzed and interpreted the data. SMS and WTW wrote and edited the manuscript. WTW supervised the study.

## Competing interests

None.

## Data and materials availability

The 661W cell line was obtained through a MTA with the Board of Regents of the University of Oklahoma.

