## Supplemental material for "Complement C3- and CR3-dependent microglial clearance protects photoreceptors in retinitis pigmentosa"

**List of Supplementary Materials**

**Supplementary Materials and Methods**

**Supplementary Figures**

Fig. S1. Complement components, regulatory factors, and receptors are not markedly increased in expression in wild type retina across the ages of P16 to P30.

Fig. S2. *C3* mRNA expression is upregulated in the ONL of histopathological retinal specimens of human retinitis pigmentosa (RP) and is localized to infiltrating microglia.

Fig. S3. C3 deficiency in the rd10 mouse model results in accelerated functional cone degeneration.

Fig. S4. Histological evidence of accelerated rod photoreceptor degeneration in C3 deficiency in the rd10 mouse model.

Fig. S5. Effect of C3 genetic deficiency in the absence of the rd10 mutation on retinal thickness and ERG function.

Fig. S6. Effect of CR3 genetic deficiency in the absence of the rd10 mutation on retinal thickness and ERG function.

Fig. S7. *In vitro* C3 opsonization of photoreceptor outer segments mediates their phagocytosis by microglia via a C3-CR3 dependent mechanism.

Fig. S8. Schematic depicting a C3-CR3-dependent mechanism of microglial phagocytic clearance of apoptotic photoreceptors in the degenerating rd10 retina.

**Supplementary Tables**

Table S1. Primers

Table S2. Antibodies

***Supplementary Materials and Methods***

***mRNA analysis by quantitative PCR***

Mice euthanized by $CO_2$ inhalation were enucleated and the retinas harvested by dissection. Cells in retina tissue were lysed by trituration and homogenized using QIAshredder spin columns (Qiagen). Total RNA was isolated using the RNeasy Mini kit (Qiagen) according to the manufacturer's specifications. First strand synthesis was performed with PrimeScript™ cDNA synthesis kit (Takara) following the manufacturer's instructions. qRT-PCR was performed using a SYBR green RT-PCR kit (Affymetrix) in CFX96 Real-Time PCR system (BioRad) under the following conditions: denaturation at 95 °C for 5 min, followed by 40 cycles of 95 °C for 10 s, and then 60 °C for 45 s. Threshold cycle (CT) values were calculated and expressed as fold-change determined using the comparative CT (2ΔΔCT) method. Housekeeping genes *Gapdh* and *Actb* were used as internal controls. Oligonucleotide primer pairs used are listed in Supplementary Table 1.

***Detection and localization of mRNA in retinal sections using multiplex in situ hybridization***

For *in situ* hybridization of mRNA probes, frozen retinal tissues from murine and human eyes were sectioned at 12µm. Resulting sections were heated at 65°C for 1 h, immersed in 4% paraformaldehyde for 15 minutes, and then in RNAscope® Protease III for 30 minutes at 40°C. *In situ* detection of mouse *C3* (Mm-C3, #417841), mouse *Cx3cr1* (Mm-Cx3cr1, #314221), and human *C3* (Hs-C3, #430701) was performed using the RNAscope® Multiplex Fluorescent Reagent Kit v2 (Advanced Cell Diagnostics) according to the manufacturer's specifications. Fluorescent signal amplification was performed with TSA® Plus fluorescein or Cy5 (1:1500, NEL760001KT, PerkinElmer).

***Immunohistochemistry and TUNEL labeling of retinal tissue***

Enucleated eyes from euthanized mice were dissected to form posterior segment eye-cups and fixed in 4% paraformaldehyde in PBS for 1-2hr at 4°C. Eyecups were either cut into 30µm-thick sections using a cryostat (Leica CM3050S) or dissected to form retinal flat-mounts. Flat-mounted retinas were blocked for 1 h in 2% Roche blocking reagent (11096176001, Millipore Sigma) in PBST at room temperature. Sections previously subjected to multiplex *in situ* hybridization were also analyzed. Primary antibodies, which included IBA1 (1:500, Wako, #019-19741), C3b/iC3b (1:100, Hycult Bio, #HM1078), CFB (1:200, Santa Cruz Bio, #sc-67141), rhodopsin (1:100, EMD Millipore, #MAB5356), were diluted in blocking buffer and incubated overnight for sections and 3 days for flat-mounts at 4°C with gentle shaking. After washing in 1X PBST, retinal samples were incubated for 1h at room temperature for sections or overnight for flat-mounts with secondary antibodies (AlexaFluor 488-, 568-conjugated anti-rabbit or mouse IgG, Invitrogen) and mounted on slides with Prolong Gold with DAPI (Invitrogen) to label cell nuclei. Apoptosis of retinal cells was assayed using a TUNEL assay (*in situ* cell death detection kit, TMR red; Roche) according to the manufacturer's specifications. Stained retinal samples were imaged with confocal microscopy (Zeiss LSM 700 or Zeiss LSM 880). For analysis at high magnification, multiplane z-series were collected using 20× or 40× objective; each z-series spanned from the outer plexiform layer (OPL) to the sub-retinal space (SRS) for retinal flat-mounts, and over a depth of 30µm for retinal sections, with each section spaced 1µm apart. Confocal image stacks were viewed and analyzed Zen software (Zeiss), and/or ImageJ (NIH).

***Histological image analysis***

Quantitative histological assessments of labeled retinal sections were used to assess the extent of photoreceptor degeneration in experimental mice. Mean outer nuclear layer (ONL) thickness was quantified by calculating the area of the ONL in an imaging field as revealed by nuclear labeling by DAPI and/or rhodopsin-labeling of rods and dividing by the length of the field. The density of apoptotic photoreceptors was calculated by manually counting $TUNEL^+$ cells in the ONL within the imaging field and dividing by the area of the ONL.

***Electroretinographic (ERG) analysis***

ERGs were recorded using an Espion E2 system (Diagnosys). Mice were anesthetized as described above after dark adaptation overnight. Pupils were dilated and a drop of proparacaine hydrochloride (0.5%; Alcon) was applied on cornea for topical anesthesia. Flash ERG recordings were obtained simultaneously from both eyes with gold wire loop electrodes, with the reference electrode placed in the mouth and the ground subdermal electrode at the tail. ERG responses were obtained at increasing light intensities over the ranges of $1X10^{-4}$ to 10 $cd/s/m^2$ under dark-adapted conditions and 0.3 to 100 $cd/s/m^2$ under a rod-saturating background light. The stimulus interval between flashes varied from 5s at the lowest stimulus to 60s at the highest ones. Two to 10 responses were averaged depending on flash intensity. ERG signals were recorded with 0.3 Hz low-frequency and 300 Hz high-frequency cutoffs sampled at 1 kHz. Analysis of a-wave and b-wave amplitudes was performed using Espion ERG Data Analyzer software (version 6.0.54). The a-wave amplitude was measured from the baseline to the negative peak and the b-wave was measured from the a-wave trough to the maximum positive peak. Statistical comparisons between ERG amplitudes between animals of different genotypes were analyzed using a two-way ANOVA.

***Image analysis of microglial features***

Microglia characteristics were assessed in immunolabelled retinal flat mount preparations and frozen sections. For each flat mount of the entire retina, 20X and 40X images were acquired in each of the four quadrants; for each quadrant, one central (located midway between the optic nerve and the equator of the globe) and one peripheral imaging field (located midway between the equator of the globe and the peripheral retinal edge) was obtained. Quantitative image analysis for infiltrating microglia number, mean microglial soma size, microglial phagosome number, and microglia number containing TUNEL+ and TUNEL- ONL nuclei, were performed manually using computer-assisted software (Analyze Particles function, NIH ImageJ). Internalization of rhodopsin by IBA1+ microglia was quantified in Imaris (Bitplane) by setting a conserved threshold for both rhodopsin and IBA1 labeling channels and computing the number of colocalized voxels.

***mRNA profiling in retinal tissue using Nanostring***

mRNA expression in retinal tissue was profiled and analyzed using the Nanostring platform nCounter Mouse Neuroinflammation panel containing 757 neuroinflammatory-related mouse genes and 13 internal reference controls (Nanostring, Seattle, WA, #115000237). Briefly, the total RNA from a single retina was extracted using the RNeasy kit (Qiagen). A total of 100ng RNA in a volume of 5µl was then hybridized to the capture and reporter probe sets at 65°C for 16 h according to the manufacturer's instructions. The individual hybridization reactions were washed and eluted per protocol at a NIH Core Facility (CCR Genomics Core, NCI) and the data collected using the nCounter Digital Analyzer (Nanostring). Generated data was evaluated using internal QC process and the resulting data were normalized with the geometric mean of the housekeeping genes using the nSolver 4.0 and Advanced Analysis 2.0 software (Nanostring). Retinas from $C3^{+/+}$; $CR3^{+/+}$, $C3^{-/-}$, and $CR3^{-/-}$ genotypes of animals in the rd10 background at time points of P24 and P30, each comprising 3-4 biological repeats, were analyzed. Differentially expressed genes were defined as those demonstrating a difference in expression level of fold change of ≥ 1.5, with a p-value of $<0.05$ (adjusted p-value, t- test). Unsupervised hierarchical clustering and heat map analysis were performed using nSolver software. Canonical pathway analyses were performed using Ingenuity Pathway Analysis (IPA, Qiagen).

***Isolation and culture of primary microglia***

Retinal microglia were isolated from P24 $C3^{+/+}$; $C3^{+/+}$.rd10, $C3^{-/-}$.rd10, and $CR3^{-/-}$.rd10 mice. Enucleated globes were immersed in ice-cold Hank's balanced salt solution (HBSS) and retinas were isolated by dissection before transfer into 0.2% papain solution including glucose (1 mg/mL), DNAse1 (100 U/mL; Worthington), superoxide dismutase (SOD) (5 mg/mL; Worthington), gentamycin (1 µL/mL; Sigma), and catalase (5 mg/mL; Sigma) in HBSS, and incubated at 8°C for 45 minutes and then at 28°C for 7 minutes. The digested tissue was dissociated by trituration and centrifuged at 150G for 5 minutes at 4°C. The resulting cell pellet was resuspended with neutralization buffer containing glucose (2 mg/mL), DNAse1 (100 U/mL), SOD (5 mg/mL), catalase (5 mg/mL), antipain (50 mg/mL; Roche), d-a-tocopheryl acetate (10 mg/mL; Sigma), albumin (40 mg/mL), and gentamycin (1 mL/mL, Sigma), and again centrifuged at 150G for 5 minutes at 4°C. The resulting pellet was resuspended in Dulbecco's Modified Eagle Medium (DMEM)/ Nutrient Mixture F-12 (1:1) media with 10% fetal bovine serum (Gibco) and 1X minimum essential medium (MEM) nonessential amino acids solution (Sigma) before transfer into 75 $cm^2$ flasks. The resulting mixed cell cultures that adhered to the bottom of the flasks were allowed to grow to confluence. Microglial cells were detached from the adherent cell layer by shaking, collected, and transferred to a new 12-well plate containing sterile glass coverslips. The culture medium was collected every 3 days and replaced with fresh culture medium. Experimentation was conducted when isolated microglia attained 40-50% confluence.

***In vitro assay for microglial phagocytosis of photoreceptor outer segments***

Microglial phagocytosis of bovine photoreceptor rod outer segments (POS) was evaluated and compared between retinal microglia isolated from mice of different genotypes. Bovine POS preparations (InVision Bioresources) were diluted in serum-free DMEM/F12 (1:1, Gibco) to a concentration of $10^6$ segments/mL and fluorescently labeled with the lipophilic dye DiI (Vybrant Cell-Labeling Solutions, Invitrogen) according to the manufacturer's instructions. Labelled POS were incubated in either 20% complete mouse serum or C3-depleted mouse serum for 1 hour at 37°C and then transferred back into serum-free media. The resulting POS suspension ($10^5$ segments in 100μl) was then added to cultured retinal microglia (50% confluency, grown on coverslips in a 12-well plate in 1ml of DMEM/F12) incubated at 37°C for 2 hours to allow for phagocytosis. Non-phagocytosed POS were removed by washing and microglia on the coverslips were fixed in 4% PFA, stained with DAPI to label nuclei, and then mounted and imaged with a Zeiss LSM 880 confocal microscope. The total number of microglia per field and the proportion of microglia incorporating fluorescently-labeled POS was measured.

***In vitro assay for microglial neurotoxicity to photoreceptors***

661W photoreceptor cells (gifted from Dr. Muayyad Al-Ubaidi) were maintained in 96-well plates. Cultured retinal microglia from animals of different genotypes were allowed to condition serum-free media (DMEM/F12, Gibco) for 48 hours. Conditioned media (100μl) was then added to 661W cells for 16 hr and the resulting cell viability assessed using an MTT cell assay kits (ATCC; Manassas, VA) following the manufacturer's specifications. Protein analysis of conditioned media was performed using a customized ProcartaPlex Multiplex Immunoassay panel (ThermoFisher) according to the manufacturer's instructions. In brief, magnetic beads were seeded in triplicate onto a 96-well plate, after which conditioned media (50μl) from retinal microglia cultured from C57BL6/J, $C3^{+/+}$; $C3^{+/+}$.rd10, $C3^{-/-}$.rd10, and $CR3^{-/-}$.rd10 animals were added to each well and incubated overnight at 4°C. The beads were washed before the addition of the detecting antibodies followed by Streptavidin-PE addition and quantification on a LUMINEX® 200™ instrument (Luminex).

### Supplementary Figures

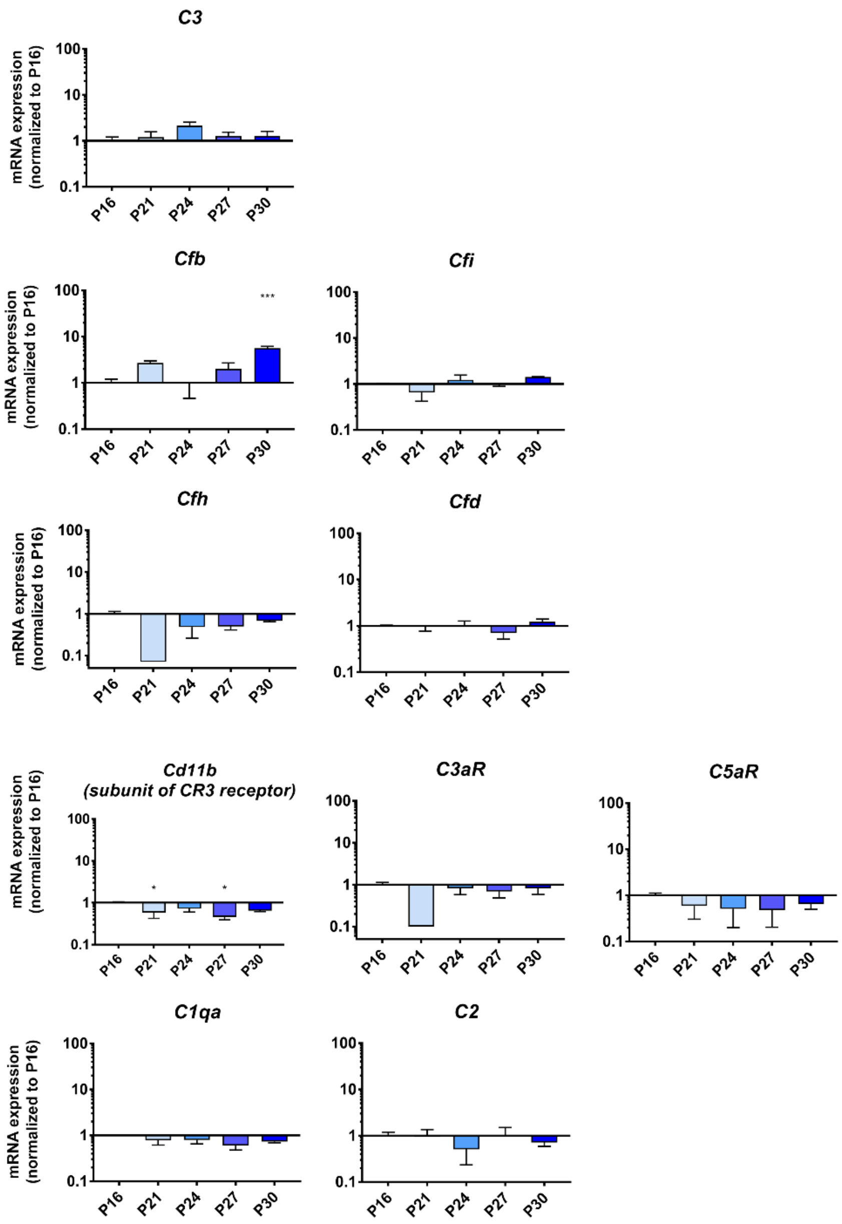


**Fig. S1. Complement components, regulatory factors, and receptors are not markedly increased in expression in wild type retina across the ages of P16 to P30.** mRNA expression levels of different complement components in wild type retina analyzed using RT-PCR did not demonstrate patterns of prominent and sustained upregulation comparable to those observed in rd10 retina during photoreceptor degeneration (P16 to P30). Expression levels for each gene at different time-points were normalized relative to that at P16, p values for comparisons relative to levels at P16 were indicated as: *, $p<0.05$; **, $p<0.01$; ***, $p<0.001$; 1-way ANOVA with Dunnett's multiple comparison test, n = 4 animals per time-point.

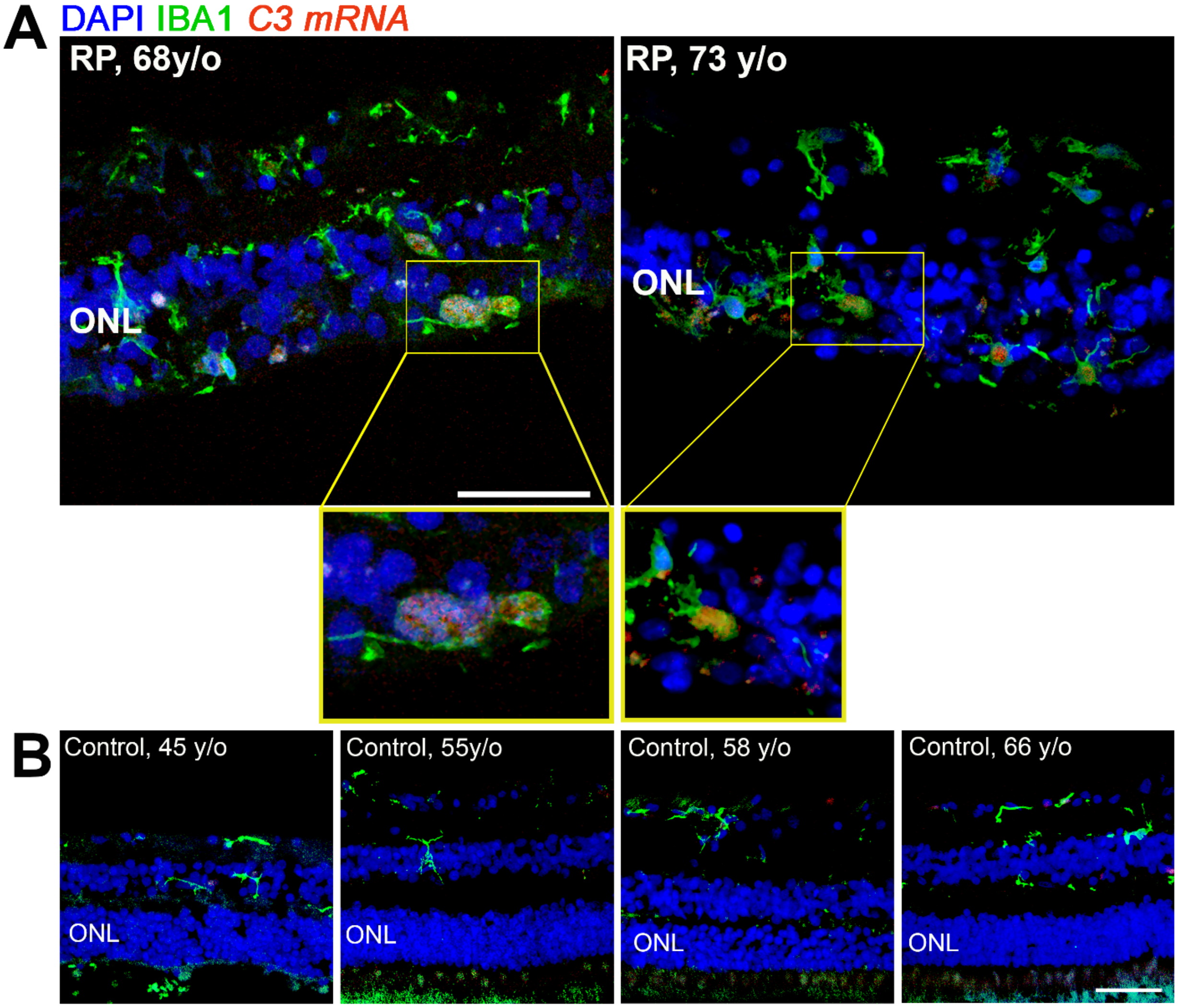


**Fig. S2. *C3* mRNA expression is upregulated in the ONL of histopathological retinal specimens of human retinitis pigmentosa (RP) and is localized to infiltrating microglia.** **(A)** Localization of *C3* mRNA expression by *in situ* hybridization (*red*) in histopathological retinal sections from two patients with RP demonstrates labeling in IBA1-immunopositive microglia that have infiltrated the ONL; insets show high-magnification views of infiltrating microglia with labeling for *C3* mRNA within DAPI-labeled nuclei. (**B**) Similar evaluations of retinal sections from healthy middle-aged control patients without a diagnosis of retinal disease (n = 4) demonstrated an absence of infiltrating microglia in the ONL or detectable *C3* mRNA expression. Scale bars = 50µm.

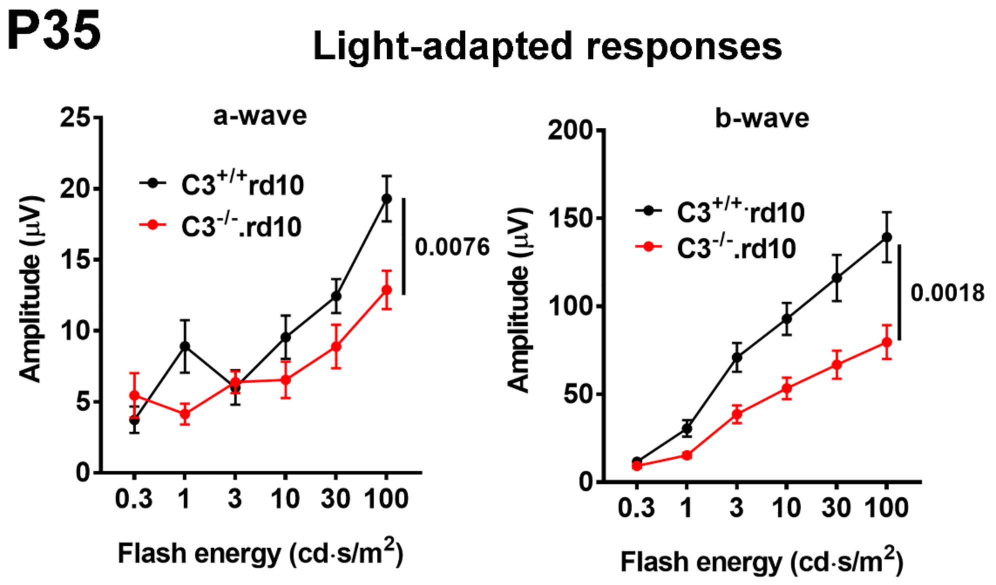


**Fig. S3. C3 deficiency in the rd10 mouse model results in accelerated functional cone degeneration.** ERG evaluation at P35 at a later time-point where cone degeneration is underway demonstrated decreased light-adapted, cone a- and b-wave amplitudes in $C3^{-/-}$.rd10 relative to $C3^{+/+}$.rd10 animals (p values were derived from a 2-way ANOVA with Tukey's multiple comparisons test, number of eyes analyzed were: $C3^{+/+}$.rd10 = 16; $C3^{-/-}$.rd10 = 14).

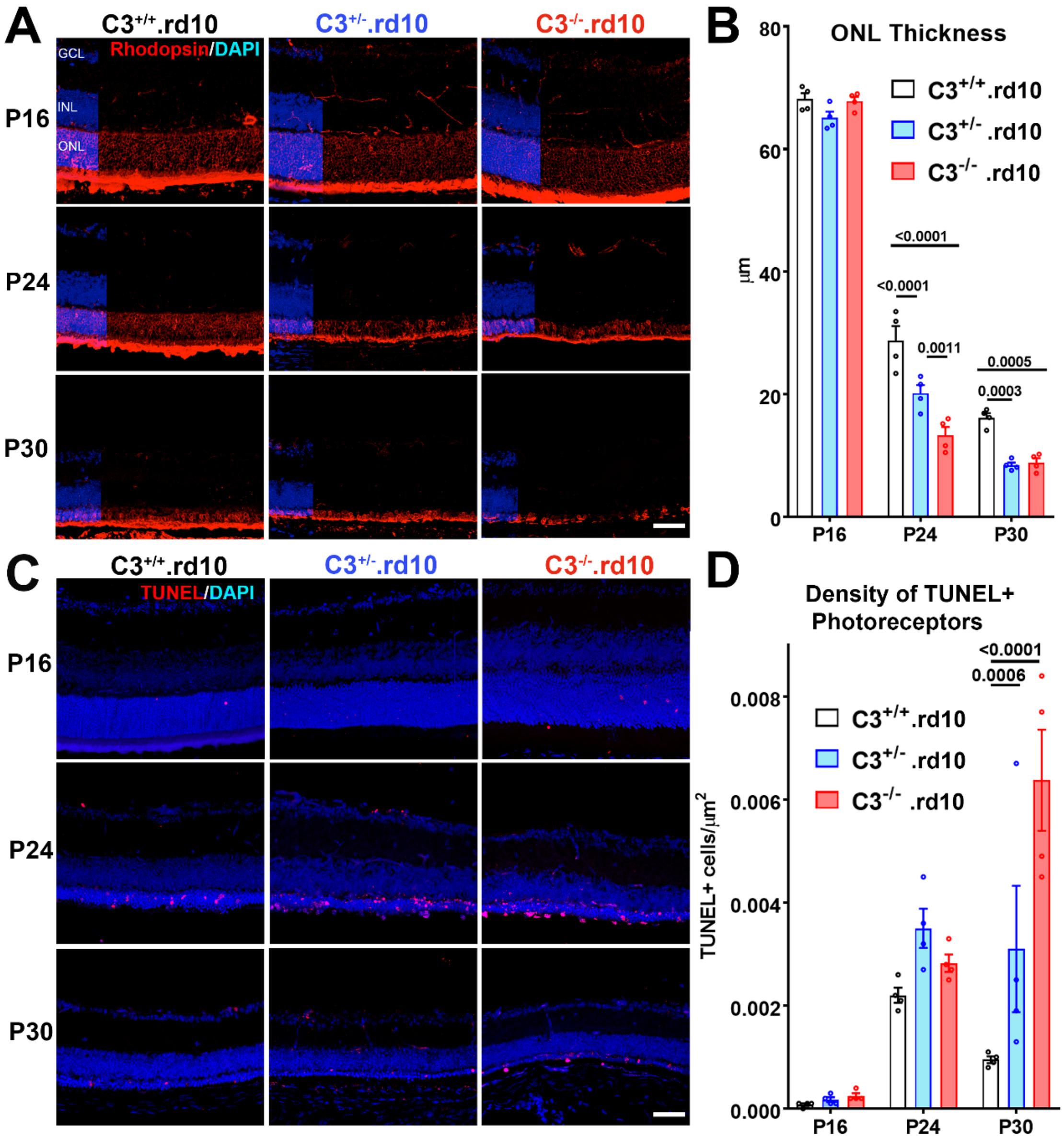


**Fig. S4. Histological evidence of accelerated rod photoreceptor degeneration in C3 deficiency in the rd10 mouse model.** (**A, B**) The effects of C3 genotype on ONL thickness and rod photoreceptor degeneration were analyzed in retinal sections from $C3^{+/+}$.rd10, $C3^{+/-}$.rd10, and $C3^{-/-}$.rd10 animals at P16, P24, and P30. Retinal lamination and ONL thickness, as revealed by nuclear labeling with DAPI (*blue*), were similar between genotypes at P16. At P24 and P30, decreases in the thickness of the ONL layer were significantly greater in $C3^{+/-}$.rd10 and $C3^{-/-}$.rd10, relative to $C3^{+/+}$.rd10 animals. Immunohistochemical analysis of rhodopsin labeling (*red*) also demonstrated more severe shortening of rod outer segments and loss of rods in $C3^{+/-}$.rd10 and $C3^{-/-}$.rd10 animals. (**C, D**) Analysis of the density of TUNEL+ apoptotic cells in the ONL showed minimal TUNEL labeling at P16 but significantly greater density in $C3^{+/-}$.rd10 and $C3^{-/-}$.rd10 animals relative to $C3^{+/+}$.rd10 animals. p values were derived from a 2-way ANOVA with Tukey's multiple comparisons test, n = 4 animal per genotype and age group. Scale bars = 50µm.

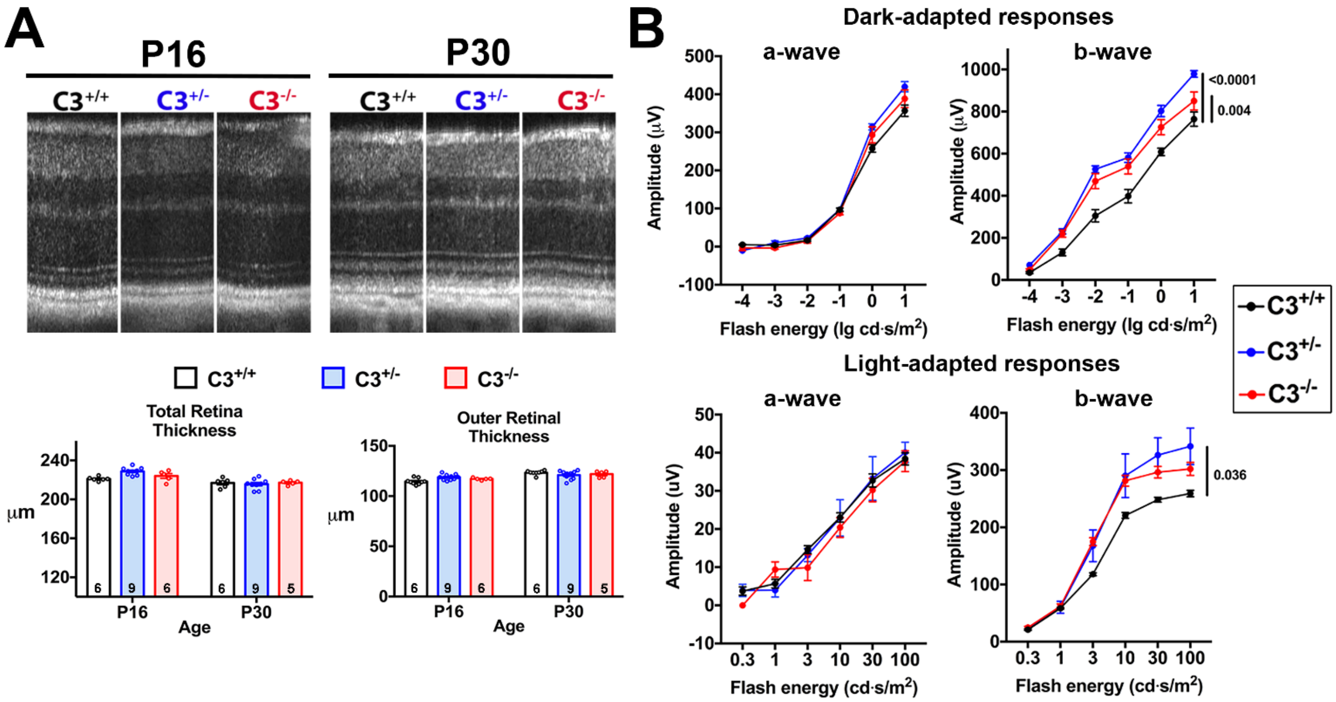


**Fig. S5. Effect of C3 genetic deficiency in the absence of the rd10 mutation on retinal thickness and ERG function**. (**A**) Retinal lamination and thickness of $C3^{+/+}$, $C3^{+/-}$, and $C3^{-/-}$ animals (in the absence of the rd10 mutation) were evaluated at P16 and P30 using *in vivo* OCT imaging. The nature of retinal lamination was similar between the 3 genotypes at both ages, and the total and outer retinal thicknesses were also statistically non-distinct. p values were derived from a 2-way ANOVA with Tukey's multiple comparisons test, number of eyes analyzed in each group is provided at the bottom of each column. (**B**) ERG evaluation of retinal function at P24 showed no significant differences between the 3 genotypes with respect to dark- and light-adapted a-wave amplitudes, indicating similarities in both rod and cone photoreceptor functions. Dark- and light-adapted b-wave amplitudes were however slightly and significantly higher in $C3^{+/-}$ and $C3^{-/-}$ animals relative to $C3^{+/+}$ animals, indicating a deficit in synaptic elimination and refinement at the level of the OPL in C3 deficiency. p values were derived from a 2-way ANOVA with Tukey's multiple comparisons test, number of eyes analyzed were: 7, 5, and 5, for $C3^{+/+}$.rd10, $C3^{+/-}$.rd10, and $C3^{-/-}$.rd10 animals respectively.

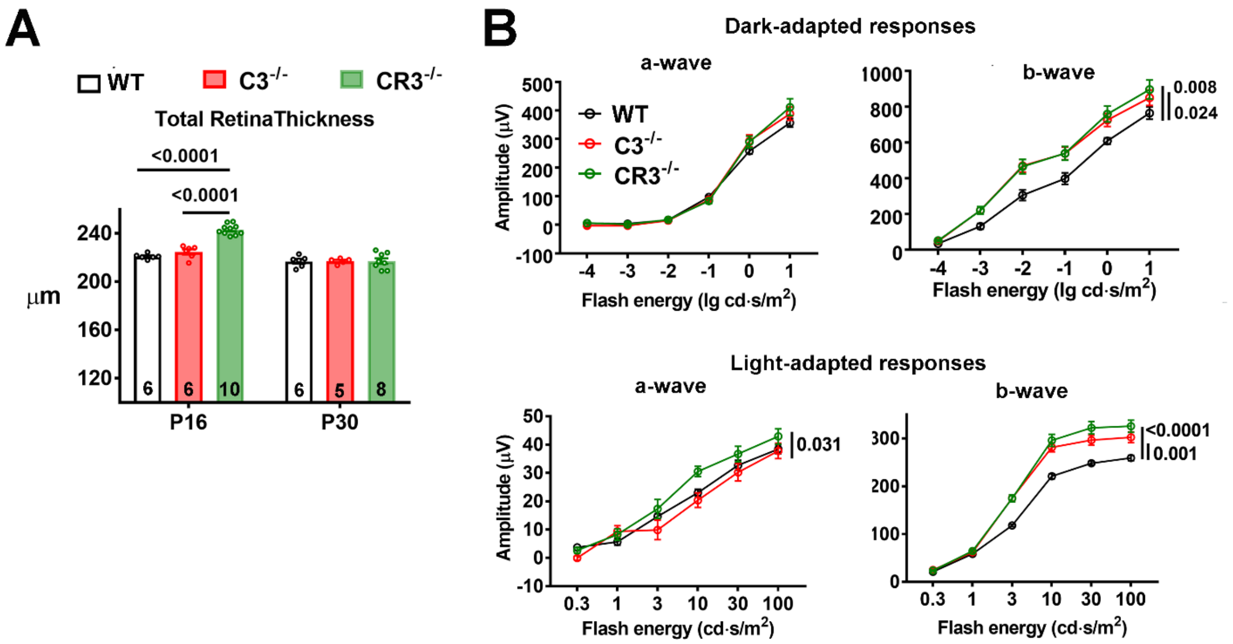


**Fig. S6. Effect of CR3 genetic deficiency in the absence of the rd10 mutation on retinal thickness and ERG function**. (**A**) Retinal thickness of WT, C3$^{-/-}$, and CR3$^{-/-}$ animals in the absence of the rd10 mutation were compared at P16 and P30 using *in vivo* OCT imaging. Total retinal thicknesses were greater in CR3$^{-/-}$ relative to the other two genotypes at P16 but were statistically similar between all 3 genotypes at P30. p values were derived from a 2-way ANOVA with Tukey's multiple comparisons test, number of eyes analyzed in each group is provided at the bottom of each column. (**B**) ERG evaluation of dark-adapted retinal function at P24 showed that CR3$^{-/-}$ animals showed similar a-wave amplitudes but increased b-wave amplitudes relative to WT animals, matching those in C3$^{-/-}$ animals. Light-adapted ERG evaluation showed a slight a-wave amplitude increase in CR3$^{-/-}$ animals relative to the other 2 genotypes, while b-wave amplitudes in CR3$^{-/-}$ animals were higher than those in WT animals, matching those found in C3$^{-/-}$ animals. p values were derived from a 2-way ANOVA with Tukey's multiple comparisons test, number of eyes analyzed were: 7, 5, and 6, for WT, C3$^{-/-}$, and CR3$^{-/-}$ animals respectively.

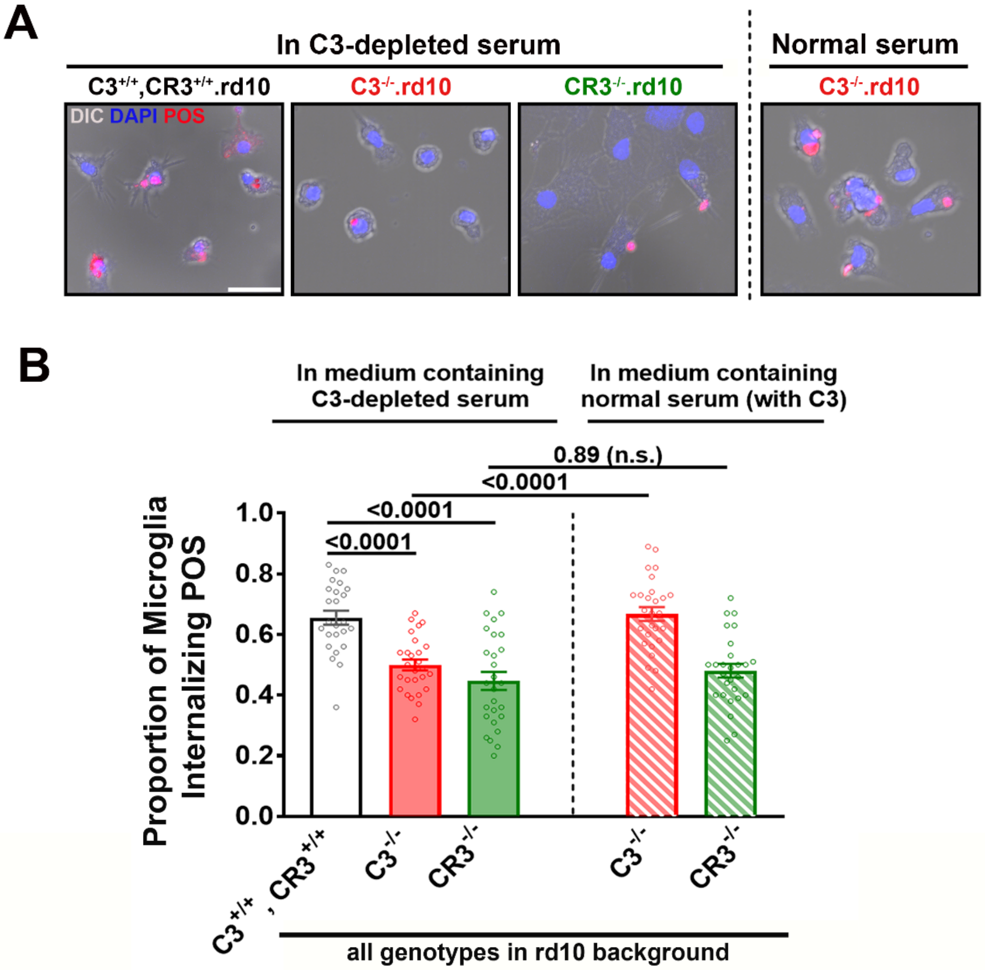


**Fig. S7.** ***In vitro*** **C3 opsonization of photoreceptor outer segments mediates their phagocytosis by microglia via a C3-CR3 dependent mechanism.** Phagocytic capacities of microglia isolated from P24 $C3^{+/+}$,$CR3^{+/+}$.rd10, $C3^{-/-}$.rd10, and $CR3^{-/-}$.rd10 retinas were evaluated in an *in vitro* phagocytic assay that assessed the phagocytic uptake of bovine rod photoreceptor outer segments (POS) added in culture. **(A)** Fluorescently-labeled photoreceptor outer segments (*red*) were incubated with cultured microglia (seen in DIC images) in the presence of either in normal, C3-sufficient or C3-depleted serum and cultured for two hours to permit microglial phagocytosis. Scale bar = 25µm. **(B)** Under C3-depleted conditions, the proportions of $C3^{-/-}$.rd10 and $CR3^{-/-}$.rd10 microglia phagocytosing POS were similar and significantly lower than that in $C3^{+/+}$;$CR3^{+/+}$.rd10 microglia, indicating a requirement for microglial production of C3 and expression of CR3 for POS phagocytosis. However, in a medium containing extracellular C3, the proportion of POS+ $C3^{-/-}$.rd10 microglia was significantly increased to the level demonstrated by $C3^{+/+}$;$CR3^{+/+}$.rd10 microglia, indicating that exogenous C3 can compensate for the absence of microglia-derived C3. Exogenous C3 did not increase the phagocytic rate in $CR3^{-/-}$.rd10 microglia, indicating a maintained requirement for microglial CR3 function. p values were derived from a 1-way ANOVA with Tukey's multiple comparisons test, n = 24-26 replicates per conditions from two independent trials.

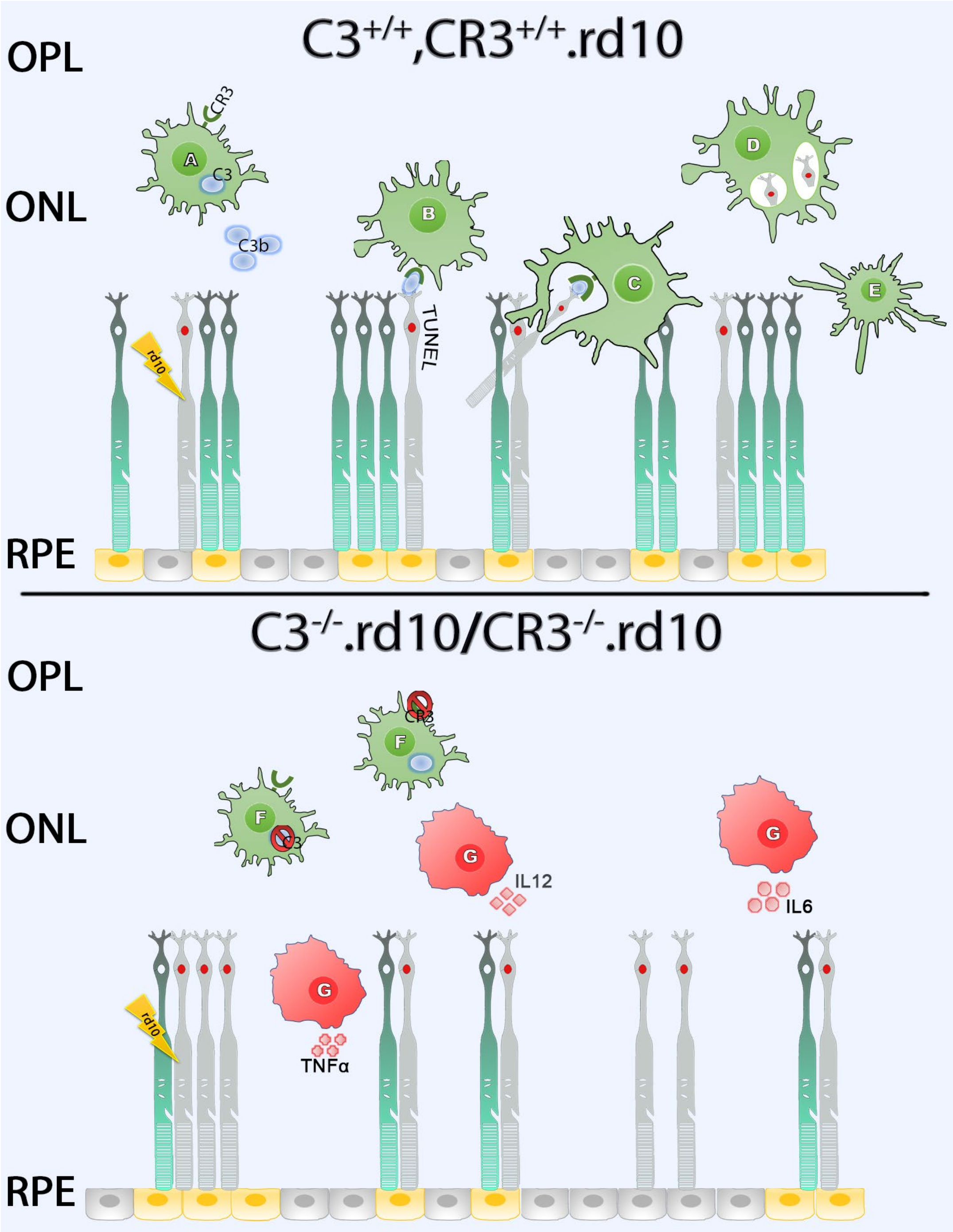


**Fig. S8. Schematic depicting a C3-CR3-dependent mechanism of microglial phagocytic clearance of apoptotic photoreceptors in the degenerating rd10 retina (*Upper panel*)** In the complement-sufficient rd10 ($C3^{+/+}$, $CR3^{+/+}$.rd10) mouse retina, microglia, sensing the degeneration of mutation-bearing rod photoreceptors, infiltrate the ONL, upregulating their expression and secretion of C3 (depicted as *blue circles*)(**A**). Extracellular activation of C3 to form iC3b results in the opsonization of apoptotic rod photoreceptors, which show TUNEL labeling (depicted in *red*)(B). CR3-expressing infiltrating microglia recognize the opsonized targets via iC3b-CR3 binding, and clear them via phagocytosis (**C**), forming phagosomes containing apoptotic rods (**D**). Following phagocytosis of apoptotic rods, infiltrating microglia are decreased in their activation status (**E**), demonstrating downregulated proinflammatory cytokine expression and decreased neurotoxic potential. (***Lower panel***) In rd10 retinas for which either C3 ($C3^{-/-}$.rd10) or CR3 ($CR3^{-/-}$.rd10) is genetically deficient (**F**), the opsonization of apoptotic rod photoreceptors or the recognition of opsonized targets is impaired respectively. Failure of microglial phagocytic clearance results in the accumulation of apoptotic TUNEL+ rods that induce a more rapid, non-cell-autonomous degeneration of nearby rod photoreceptors. This potentiation in degeneration may be driven by an increased activation of non-phagocytic microglia (depicted as *red cells*)(**G**), that may result from increased stimulation from signals (e.g. danger-associated molecular patterns) released from apoptotic cells, or decreased downregulatory effects that typically follow microglial phagocytosis of apoptotic cells. These together may increase the secretion of pro-inflammatory cytokines (TNFα, IL6, IL12), augmenting microglial neurotoxicity to photoreceptors, and accelerating the course of structural and functional degeneration in the rd10 retina.

**Supplemental Tables**

**Table S1. List of primers used for RT-PCR reactions**

| Gene | Forward | Reverse |
|---|---|---|
| **C1qa** | CAAGGACTGAAGGGCGTGAA | CAAGCGTCATTGGGTTCTGC |
| **C2** | CTCATCCGCGTTTACTCCAT | TGTTCTGTTCGATGCTCAGG |
| **C3** | GAAGTACCTCATGTGGGGCC | CAGTTGGGACAACCATAAACC |
| **C3aR** | GGAAGCTGTGATGTCCTGG | CACACATCTGTACTCATATTGT |
| **C5aR** | GAGGGTGGAGAAGCTGAAC | CTACACCGCCTGACTCTTC |
| **Cfb** | GAGGATGGGCACAGCCCAG | GACCATATCGTGGCCTCACC |
| **Cfd** | GCAGAGAGCAACCGCAGG | CAGGATGTCATGTTACCATTTG |
| **Cfh** | CTTACATGCATGTGTAATACCA | TTATACACAAGTGGGATAATTGA |
| **Cfi** | CCATTGATGCCTGCAAAGGA | CAGACATTGTGTTGAGAAACAA |
| **CD11b** | GCAGGAGTCGTATGTGAGG | TTACTGAGGTGGGGCGTCT |

**Table S2. List of antibodies used for immunohistochemical analyses**

| Antibody | Company | Catalog # | Dilution |
|---|---|---|---|
| **C3b/iC3b** | Hycult Bio | HM1078 | 1:100 |
| **Cfb** | Santa Cruz Bio | sc-67141 | 1:200 |
| **Iba1** | Wako | 019-19741 | 1:500 |
| **Rhodopsin** | EMD Milipore | MAB5356 | 1:200 |
| **Alexa 488** | Thermo Fisher | | 1:250 |
| **Alexa 568** | Thermo Fisher | | 1:250 |
